# Enzyme Activity Regulates Substrate Diffusion by Modulating Viscosity in Crowded Milieu

**DOI:** 10.1101/2024.09.28.615560

**Authors:** Alessandro Bevilacqua, Mauricio Rios Maciel, Mark V. Sullivan, Stefano Pascarelli, Mirco Dindo, Amy Q. Shen, Paola Laurino

## Abstract

Enzymatic activity and its tight regulation are fundamental to life, yet how enzymes maintain efficient substrate availability within the highly crowded, heterogeneous native cytosol remains poorly understood. Here, we harness liquid-liquid phase separation (LLPS) to generate controlled *in vitro* protein droplets that mimic the crowding of cytosolic proteins and demonstrate that enzymatic activity enhances substrate diffusion within the crowded environment by up to three-fold. Using fluorescence microscopy, fluorescence recovery after photobleaching (FRAP), molecular docking, and particle-tracking microrheology, we show that this enhanced diffusion arises from a reduction in apparent shear viscosity driven by enzymatic disruption of transient non-covalent interactions between substrate and protein crowder. Our findings reveal a physical feedback mechanism by which enzymatic activity actively reshapes its local environment to sustain substrate availability, providing new insight into how enzymatic activity creates a positive physical feedback on its own substrate supply under the diffusion constraints of the crowded cell.

## Introduction

The physical properties of the cytosol, particularly viscosity and macromolecular crowding, are fundamental organisational principles governing cellular function, influencing macro-molecule assembly,^1^ metabolite diffusion rates,^2,3^ and cellular sensing.^4,5^ Cells dynamically modulate these properties through mechanisms such as the liquid-to-glass transition,^6–8^ changes in viscosity,^9–11^ and active viscoadaptation. ^12,13^ These discoveries have established that the physical state of the intracellular environment is dynamically regulated and, in turn, plays a central role in controlling molecular diffusion and biochemical activity. However, how such physical regulation influences substrate diffusion and enzymatic function remains incompletely understood.

Enzymes are essential for life and their activity must be precisely regulated, typically through mechanisms that modulate enzyme abundance or catalytic efficiency. ^14–22^ Extensive work using liquid-liquid phase separation (LLPS) system, has confirmed that the physical properties of the crowded medium profoundly influence enzyme activity by altering substrate diffusion and enzyme dynamics.^23–37^ Recent experiments have further shown that the reverse is also true: enzymatic activity can fluidise the crowded condensate environment, as evidenced by increased diffusion of crowder proteins and tracer particles in active versus inactive droplets.^38^ However, the molecular mechanism underlying this viscosity reduction and its direct consequences for substrate diffusion remain unresolved. Addressing this requires both direct rheological measurements and a systematic link between molecular substrate-crowder interactions and macroscopic viscosity changes.

In vitro Liquid-liquid phase separation (LLPS) systems offer a valuable tool to study these phenomena, and have been utilized to study enzyme kinetics,^29,39–41^ coacervate formation and nucleation, ^42–45^ cell growth,^46^ protein-membrane and protein-DNA interactions,^47–49^ molecular diffusion,^50^ enzyme-driven motion and migration.^51,52^ Despite the availability of LLPS systems that contain enzymes, studying the perturbation of their physical properties presents challenges due to LLPS poor stability.^53–55^ Nevertheless, *in vitro* LLPS systems remain a key approach to exploring how viscosity and crowding shape substrate diffusion and enzymatic function.

Here, using a PEG-BSA droplet system^40^ with *β*-galactosidase as a model enzyme, we combine particle-tracking microrheology, molecular docking, and fluorescence recovery after photobleaching (FRAP) to demonstrate that enzymatic activity enhances substrate diffusion in a crowded protein environment, and to identify the underlying mechanism and its molecular origin (Fig. 1).

**Figure 1:**
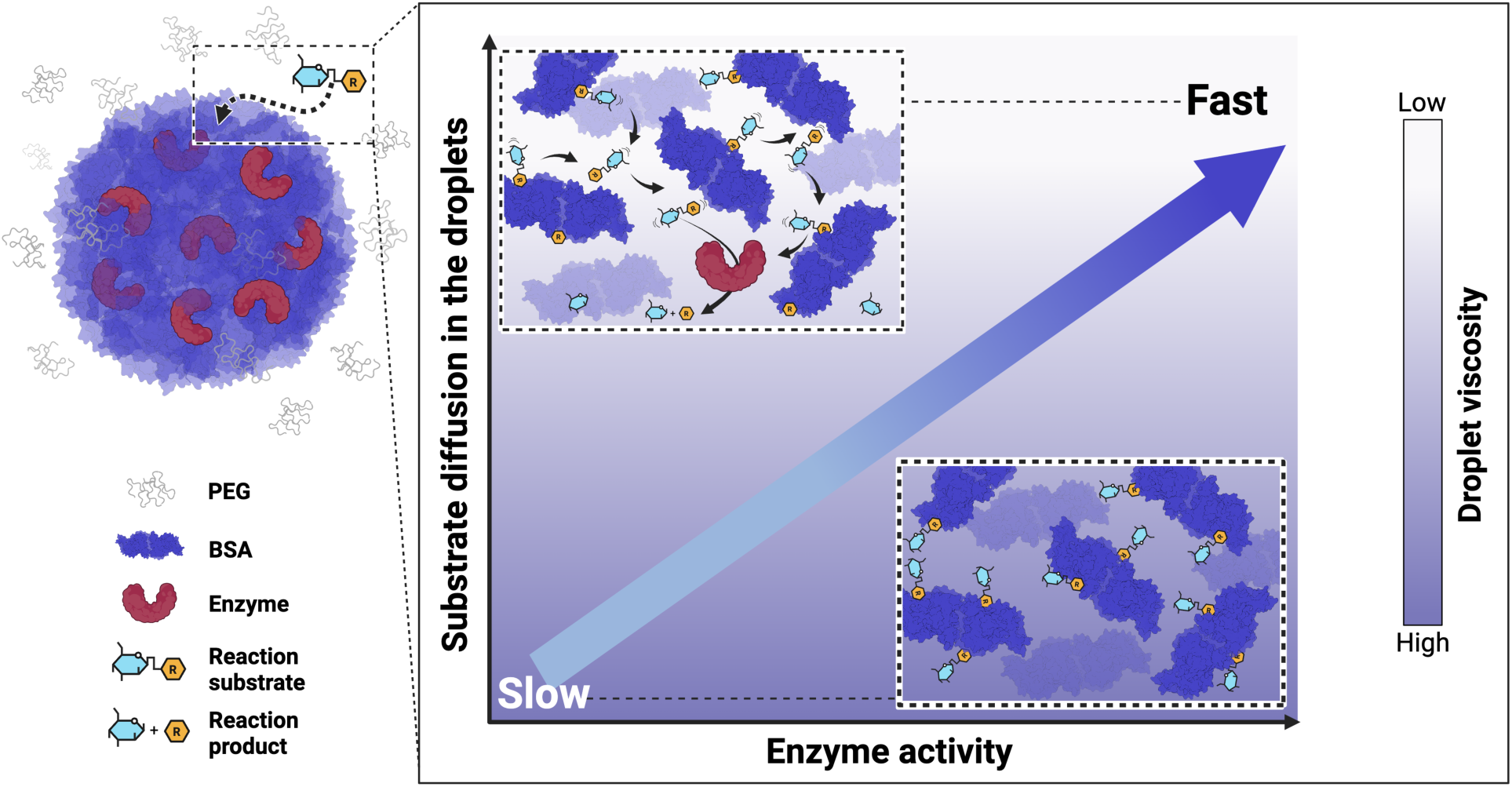
Enzymatic activity modulates substrate diffusion by decreasing the relative apparent viscosity of protein-rich droplets formed via PEG-BSA phase separation. Schematic representation of the PEG-BSA system (left), showing the formation of BSA-rich droplets that mimic cytosolic protein crowding. Enzymes preferentially partition into the droplets, while small-molecule substrates freely diffuse from the surrounding PEG-rich phase, enabling reactions within the crowded interior. The scheme plot represents the mechanism of the dynamic displacement of weak BSA-substrate bonds by which the enzymatic activity increases substrate diffusion by decreasing the relative apparent viscosity of the droplets. This image was created with BioRender.com.

We demonstrate that enzymatic turnover reduces apparent viscosity of crowded protein droplets by disrupting transient non-covalent substrate-BSA interactions and that this physical remodeling has a direct functional consequence: substrate transport within the crowded phase is enhanced by up to three-fold under diffusion-limited conditions. Molecular docking and surface plasmon resonance further reveal that reaction products interact broadly and with different affinities across the BSA surface, supporting a model in which catalytic turnover dynamically remodels transient small-molecule–crowder interactions. These findings identify enzymes as active modulators of their physical environment and provide a substrate-affinity-based mechanistic framework linking molecular interactions to diffusion in crowded biological systems.

## Results

### Enzyme kinetic characterization in the droplet system

To investigate enzyme activity in a protein-crowded environment, we utilized a PEG-BSA droplet system^40^ optimized to partition *β*-galactosidase into the BSA-rich droplets (Fig. S1). The system was characterized to determine the phase composition and droplet volume fraction under the current buffer conditions. Inside the droplets, the BSA concentration is 6.6 ± 0.5 mM, the PEG concentration is 2.8 ± 0.3 %(w/w), and the droplets comprise 3.3 ± 0.5 % of the mixture volume fraction (*ε_d_*) (Fig. S2). Assuming perfect enzyme partitioning inside the droplets (*≈* 99%, see Fig.S1), the concentration of the enzyme inside the droplets ([*E*]*_d_*) is estimated from the bulk solution concentration ([*E*]*_s_*) and the volume fraction of the droplets (*ε_d_*) as follows:

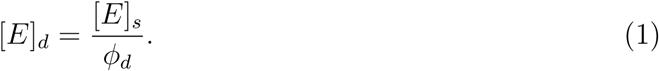

Consequently, with *ε_d_* = 3.3%, the enzyme concentration inside the droplets is therefore approximately 30-fold higher than in the bulk solution. ^40^

The enzyme activity was evaluated for the enzyme placed both within the droplets and in the PEG phase only (see Supplementary Materials and Methods). We examined the Michaelis-Menten kinetics of *β*-galactosidase with three different reaction substrates: 9H-(1,3-Dichloro-9,9-Dimethyl acridin-2-One-7-yl)-*β*-D-Galactopyranoside (DDAO-gal), D-lactose, and 4-nitrophenyl-*β*-D-galactopyranoside (ONP-gal), the reaction scheme is reported in Fig. S3 and Table S1. The kinetic parameters for *β*-galactosidase on these three substrates were derived from Michaelis-Menten kinetics (Figs. S4-S6) and are summarized in Table 1.

**Table 1:**
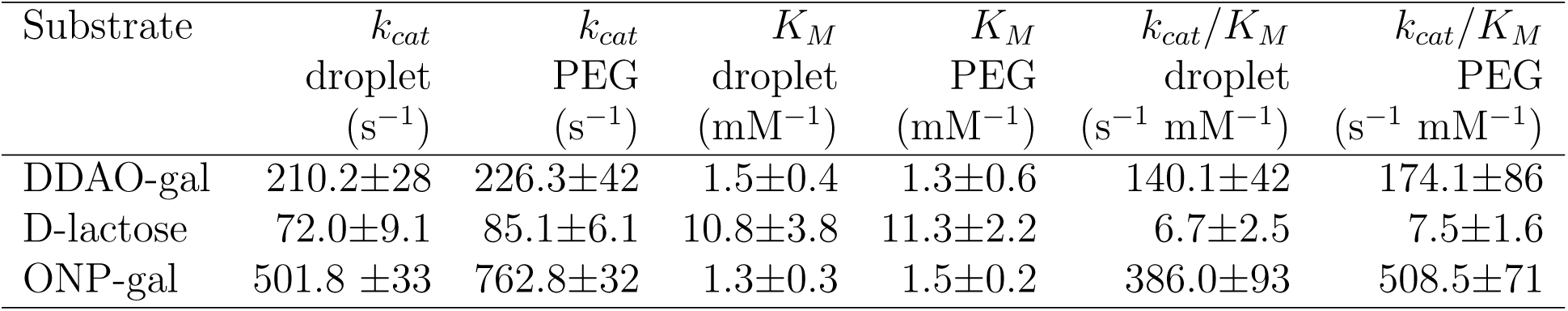
Comparison of the *β*-galactosidase kinetic parameters measured in the droplets and in the PEG-rich phase.

| Substrate | $k_{cat}$<br>droplet<br>($s^{-1}$) | $k_{cat}$<br>PEG<br>($s^{-1}$) | $K_M$<br>droplet<br>($mM^{-1}$) | $K_M$<br>PEG<br>($mM^{-1}$) | $k_{cat}/K_M$<br>droplet<br>($s^{-1} mM^{-1}$) | $k_{cat}/K_M$<br>PEG<br>($s^{-1} mM^{-1}$) |
| --- | --- | --- | --- | --- | --- | --- |
| DDAO-gal | 210.2±28 | 226.3±42 | 1.5±0.4 | 1.3±0.6 | 140.1±42 | 174.1±86 |
| D-lactose | 72.0±9.1 | 85.1±6.1 | 10.8±3.8 | 11.3±2.2 | 6.7±2.5 | 7.5±1.6 |
| ONP-gal | 501.8 ±33 | 762.8±32 | 1.3±0.3 | 1.5±0.2 | 386.0±93 | 508.5±71 |

For all three reaction substrates tested, the *K_M_* values for a given substrate do not show any significant differences when the enzyme is localized either in the BSA-rich droplets or in the surrounding PEG-phase (≤1.2-fold changes). In contrast, the enzyme shows different turnover number (*k_cat_*) for different reaction substrates in the droplet and PEG phase (Tab.1). Specifically, the enzyme compartmentalized in the droplets shows a wide range of *k_cat_* values on different substrates, representing different catalytic efficiencies (spanning from a minimum of 72.0±9.1 s^−1^ on D-lactose to 501.8 ±33 s^−1^ for ONP-gal). Inside the PEG phase the enzyme shows similar turnover numbers with a maximum 1.5-fold change in the case of ONP-gal (Tab.1).

These results establish the PEG–BSA droplets as a well-defined model system in which *β*-galactosidase remains catalytically active under crowded conditions and exhibits a broad range of turnover rates depending on the substrate. We therefore next examined whether enzymatic turnover alters droplet viscosity and whether such changes influence substrate diffusion within the crowded phase.

### Enzyme activity tunes the apparent viscosity of crowded droplets

To assess how enzymatic activity influences the microscopic rheology of the droplets environment, we combined particle-tracking microrheology and fluorescence recovery after photobleaching (FRAP), two complementary approaches that independently probe molecular mobility and local apparent viscosity (Fig. 2 A). Because FRAP recovery reflects molecular diffusion, faster recovery is consistent with lower local apparent viscosity.

**Figure 2:**
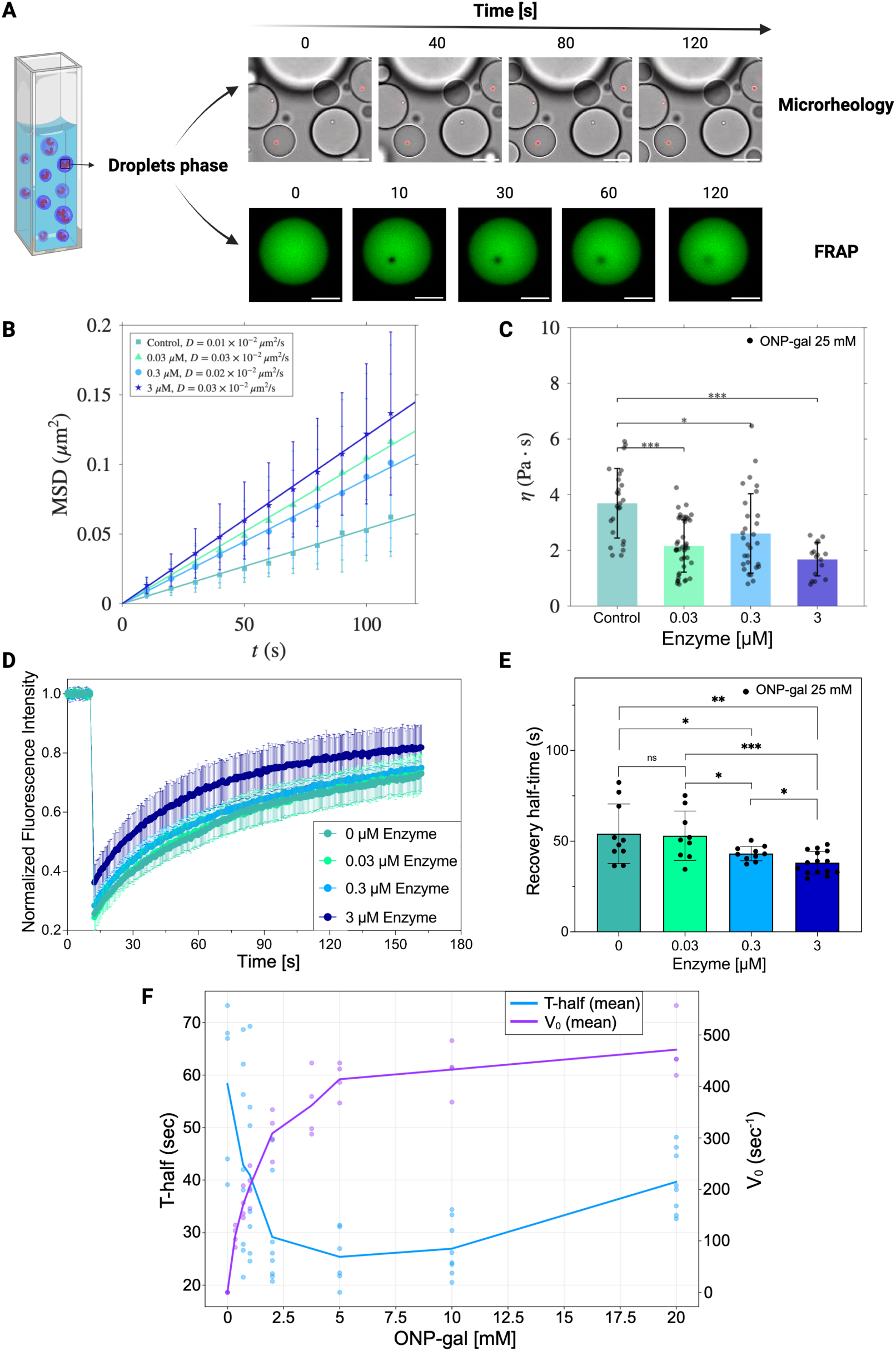
Enzymatic activity reduces the apparent viscosity of crowded protein droplets. **A** Schematic overview of the experimental setup for assessing shear viscosity within BSA-rich droplet phase using particle-tracking microrheology and fluorescence recovery after photobleaching (FRAP). Representative time-lapse images are shown for microrheology (0, 40, 80, and 120 s; scale bar: 10 *µ*m) and FRAP (0, 10, 30, 60, and 120 s; scale bar: 10 *µ*m). **B** Mean squared displacement of fluorescent tracer particles at increasing *β*-galactosidase concentrations. **C** Relative apparent viscosity, *η\**, calculated from the tracer diffusion coefficients in the presence of 25 mM ONP-gal. **D** FRAP recovery curves of Alexa Fluor 488–labeled BSA droplets following photobleaching, showing faster half-time fluorescence recovery at higher enzymatic activities (lower apparent viscosity). Each recovery curve was normalized by a reference region ROI to correct for overall photobleaching. Data represent mean ± SD (n*≥*9). **E** Relative FRAP half-recovery time (T-half) of droplets containing a range of 0–100 nM enzyme in the solution (0 - 3 *µ*M *β*-galactosidase inside the droplets) in the presence of 25 mM ONP-gal. The boxes represent the mean values and the whiskers are the SD (n*≥*9). **F** FRAP half-recovery time of droplets resembles the reaction turnover of ONP-gal. The graph shows the relative FRAP half-recovery time of droplets containing 100 nM enzyme in the solution (3 *µ*M *β*-galactosidase inside the droplets) in the presence of 0-20 mM ONP-gal and the velocity of the reaction in the presence of the corresponding substrate concentration derived from the Michaelis-Menten data of Fig.S4. All data points are shown (n*≥*4). Statistical significance was assessed using t-test. P-values are indicated for comparisons between enzyme concentrations: (0 *µ*M vs 0.03 *µ*M, p *>* 0.05; 0 *µ*M vs 0.3 *µ*M, p = 0.028; 0 *µ*M vs 3 *µ*M, p = 0.001; 0.03 *µ*M vs 0.3 *µ*M, p = 0.022; 0.03 *µ*M vs 3 *µ*M, p = 0.0007; 0.3 *µ*M vs 3 *µ*M, p = 0.016). *p *<* 0.05, **p *<* 0.01, ***p *<* 0.001. This image was created with BioRender.com.

Particle-tracking microrheology was first used to quantify changes in the local apparent viscosity of the droplet phase. Brownian trajectories of more than 50 fluorescent tracer particles (1.1 *µ*m diameter) from more than three independent experiments were recorded over 2 min for each condition (SI Movies 1–4). The diffusion coefficient, *D*, was obtained from the mean squared displacement (MSD) according to MSD(*t*) = 4*Dt*, assuming two-dimensional diffusion, and the apparent viscosity was calculated using the Stokes–Einstein equation (Materials and Methods). Measurements were performed in the presence of 25 mM ONP-gal with *β*-galactosidase concentrations of 0, 0.03, 0.3, and 3 *µ*m inside the droplets, corresponding to 0, 0.01, 0.1, and 1 nM enzyme in the bulk solution. Increasing enzymatic activity produced progressively larger MSDs (Fig. 2 B) and correspondingly lower relative apparent viscosities (Fig. 2 C), indicating that catalytic turnover fluidizes the crowded droplet phase.

To independently verify this observation using a molecular probe, we performed FRAP measurements on Alexa Fluor 488-labelled BSA, the principal structural component of the droplets. BSA was chosen because it forms the droplet-rich phase but does not participate directly in the enzymatic reaction. The BSA was labeled with the fluorescent probe Alexa Fluor 488 and used to generate droplets for the FRAP analysis (SI methods). Droplets containing the same range of enzyme concentration of the microrheology measurements (0-3 *µ*M inside the droplets) in the presence of 25 mM ONP-gal were subjected to FRAP (Fig. 2D and SI movies 5-8). The resulting recovery curves showed progressively shorter half-recovery times with increasing enzyme concentration (Fig. 2E). Specifically, in the presence of 25 mM ONP-gal the half-recovery time decreased from 54.1±16.4 in the absence of enzyme to 38.1±6.2 s at the maximum enzyme concentration tested (3 *µ*M *β*-galactosidase in the droplets), consistent with a reduction in local apparent viscosity.

Control experiments showed that neither the enzyme (0–3 *µ*M *β*-galactosidase inside the droplets) nor the substrate (0–25 mM ONP-gal) produced significant changes in the FRAP half-recovery time in the absence of enzymatic turnover (Fig. S7). The presence of the reaction product (0–25 mM ONP) results in only minimal changes in the T-half (Fig. S7). These controls demonstrate that the observed increase in molecular mobility arises specifically from catalytic activity rather than from the individual molecular components.

To determine whether the reduction in apparent viscosity depends directly on catalytic turnover, we measured the FRAP half-recovery time over a substrate concentration range of 0–20 mM ONP-gal while maintaining a constant enzyme concentration of 100 nM in the bulk solution, corresponding to 3 µM inside the droplets. As the reaction velocity increased, the FRAP half-recovery time decreased correspondingly, following a trend similar to the Michaelis–Menten dependence of the enzymatic reaction (Fig. 2F). In summary, particle-tracking microrheology and FRAP independently demonstrate that enzymatic activity reduces the apparent viscosity of the crowded droplet phase in both an enzyme-concentration- and turnover-dependent manner. Having established that catalytic turnover fluidizes the crowded protein environment, we next examined whether this physical remodeling enhances substrate diffusion within the droplets.

### Enzymatic activity enhances substrate diffusion in crowded droplets

Having established that enzymatic activity lowers the apparent viscosity of the crowded droplet phase, we next asked whether this physical remodeling enhances substrate diffusion. Although previous enzyme kinetics data are independent of the droplet size (Fig. S8), in larger droplets (diameter ≥ 40*µ*m), substrate diffusion becomes the rate-limiting step of the reaction. Here, we demonstrate how the enzyme activity can counteract these diffusion limitations by reducing the viscosity, which in turn enhances the speed of substrate diffusion.

The reaction substrate selected for this experiment is DDAO-gal since it is a fluorescent molecule that has two different excitation/emission spectra before and after the cleavage of the *β* bound between the DDAO and the D-galactose molecule^56^ by the enzyme (reaction scheme in Fig.S3). Thus, the diffusion of the substrate inside of the droplets and the generation/diffusion of the product were tracked over time by exciting the probes at *λ_g_* = 488 nm and *λ_g_* = 648 nm, respectively (Fig. 3A).

**Figure 3:**
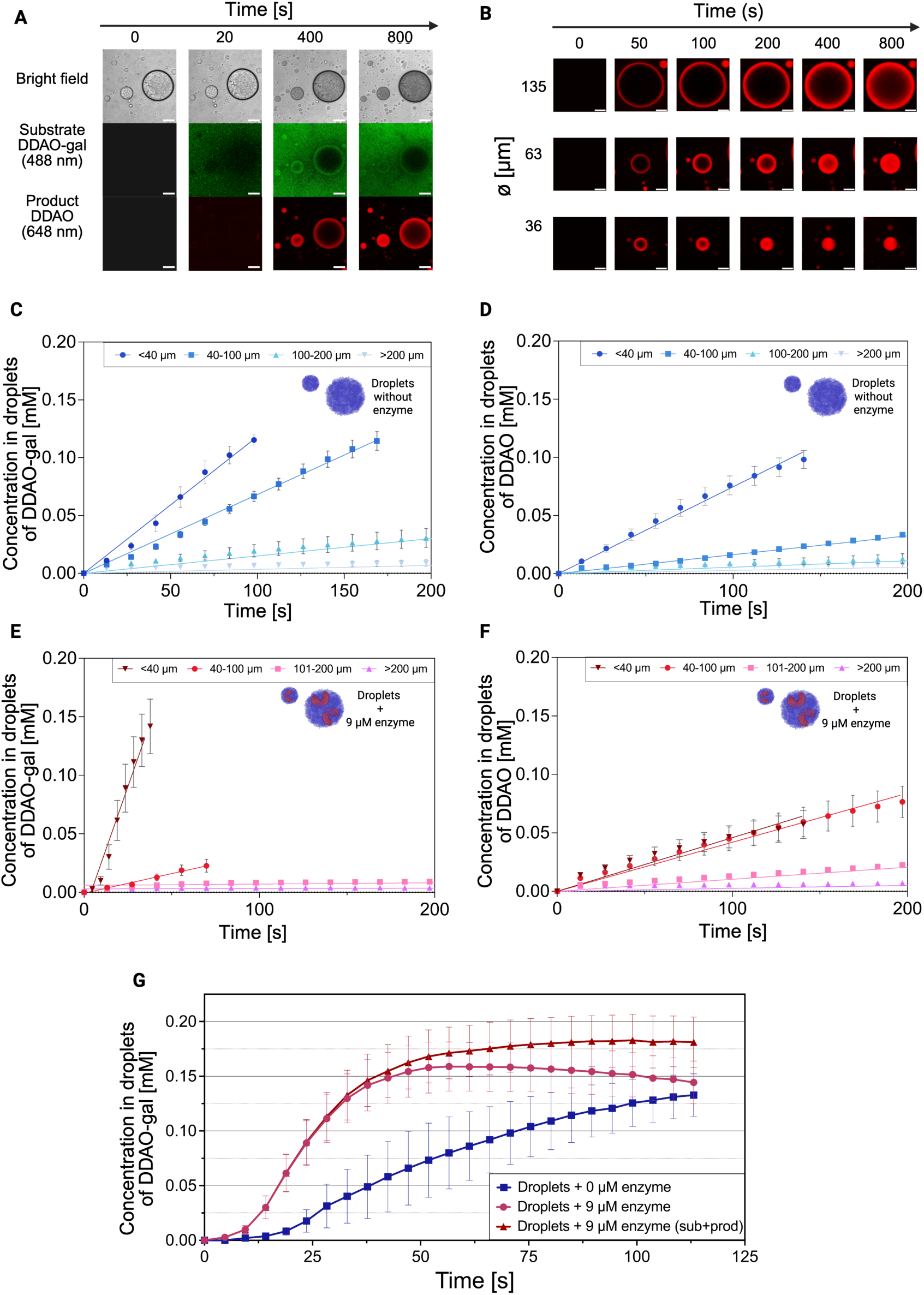
Reduction in apparent viscosity by enzymatic activity enhance substrate diffusion. **A** Visualization of the reaction occurring inside the droplets at the confocal microscope over time. Top to bottom: bright field image of the BSA droplets containing 9*µ*M *β*-galactosidase, the fluorescence of DDAO-galactoside (reaction substrate) excited at *λ_g_* = 488 nm, and the fluorescence of DDAO (reaction product) excited at *λ_g_* = 648 nm. Screenshots from SI movie 9. Scale bar 50 *µ*m. **B** Imaging of product generation (excited at 648 nm) inside droplets of different sizes (36, 63, and 135 *µ*m) over 800 seconds. Screenshots from SI movie 9. Scale bar 30 *µ*m. **C** diffusion of DDAO-Gal (reaction substrate) inside droplets (without enzyme) of a ∅ *<*40 (dark blue), 41 - 100 (blue), 101 - 200 (light blue) and *≥* 200 *µ*m (light grey). The graph shows the average concentration inside the droplets over 200 seconds from the addition of the reaction substrate. The data points at the diffusion plateau have been removed to fit the diffusion slope of the linear diffusion phase only. The full diffusion curves are reported in Fig.S10. The data represent the mean values ± SEM (n*≥*4). **D** diffusion of DDAO (reaction product) inside droplets (without enzyme) of a ∅ *<*40 (dark blue), 41 - 100 (blue), 101 - 200 (light blue) and *≥* 200 *µ*m (light grey). The graph shows the average concentration inside the droplets over 200 seconds from the addition of the reaction product. The data points at the diffusion plateau have been removed to fit the diffusion slope of the linear diffusion phase only. The full diffusion curvesare reported in Fig.S10. The data represent the mean values ± SEM (n*≥*3). **E** diffusion of DDAO-Gal (reaction substrate) inside droplets (containing 9*µ*M enzyme) of a ∅ *<* 40 (dark red), 41 - 100 (red), 101 - 200 (pink) and *≥* 200 *µ*m (light purple). The graph shows the average concentration inside the droplets over 200 seconds from the addition of the reaction substrate. The data points at the diffusion plateau have been removed to fit the diffusion slope of the linear diffusion phase only. The full diffusion curves are reported in Fig.S10. The date represents the mean values ± SEM (n*≥*5). **F** Generation of DDAO (reaction product) inside droplets (containing 9*µ*M enzyme) of a ∅ *<* 40 (dark red), 41 - 100 (red), 101 - 200 (pink) and *≥* 200 *µ*m (light purple). The graph shows the average concentration inside the droplets over 200 seconds from the addition of the reaction substrate. The data points at the diffusion plateau have been removed to fit the diffusion slope of the linear diffusion phase only. The full diffusion curves are reported in Fig.S10. The data represent the mean values ± SEM (n*≥*5). **G** Substrate diffusion is improved in enzymatically active droplets. It represents the average substrate concentration inside BSA droplets (blue) and BSA-enzyme droplets (pink) over 115 seconds. In red it is reported the sum of the average concentration of substrate and product inside BSA-enzyme droplets over 115 seconds. The droplets analyzed have a ∅ *<*40*µ*m. The data represent the mean values ± SEM (n*≥*6). This image was created with BioRender.com.

A calibration curve (Fig.S9) was established to correlate the fluorescence intensity of the substrate and product with their respective concentrations (see Supplementary Materials and Methods). Subsequently, we examined the diffusion of both substrate and product in droplets ranging in diameter (∅) from 15 to 426 *µ*m, both with and without the presence of the enzyme. A qualitative comparison of droplets containing 9*µ*M enzyme of a ∅ of 36, 63 and 135 *µ*m showed that the time needed for the reaction product DDAO to be generated in the center of the droplet increased according to the diameter of the droplet (Fig. 3B). The diffusion of substrate and product was then quantitatively evaluated by tracking the fluorescence changes inside the whole droplets over time by using a MathLab script (Fig.S10). Specifically, the movies were recorded for 1500 s from the addition of the substrate/product (0.2 mM) to reach the diffusion plateau for droplets of diameter *≤* 100*µ*m (Fig.S10). The concentration rate (here defined as the slope of the linear diffusion phase of the reaction substrate/product) in droplets without enzyme is reported in Fig. 3C-D. As expected, the concentration rate of both the substrate and product decrease by increasing the size of the droplets. This decrease reflects the reduced surface-to-volume ratio of larger droplets, making substrate diffusion increasingly diffusion-limited. In detail, the substrate concentration rate halves from droplets of ∅ *<* 40*µ*m (1.18 ± 0.19*µ*M/s) to droplets ∅ = 41- 100*µ*m (0.68 ± 0.11*µ*M/s) and keep decreasing up to 0.034 ± 0.002*µ*M/s in droplets of ∅ *>* 200*µ*m (Fig. 3C). Product diffusion exhibited a comparable size dependence (Fig. 3D).

We next repeated these measurements in droplets containing 9 *µ*M *β*-galactosidase. As expected, substrate accumulation remained increasingly diffusion-limited as droplet size increased (Fig. 3E). However, active droplets smaller than 40 *µ*m exhibited approximately threefold faster substrate accumulation than enzyme-free droplets (4.28 versus 1.18 *µ*M s1), demonstrating that enzymatic activity substantially enhances substrate diffusion (Fig. 3C, E). In contrast, larger active droplets showed lower apparent substrate accumulation than inactive droplets because the substrate was consumed by the enzymatic reaction before diffusing throughout the droplet. Consistent with this interpretation, the reduced substrate concentration observed in larger active droplets was accompanied by equivalent or higher product concentrations, compared to the enzyme-less ones (Fig. 3D, F). This fact indicates that substrate consumption, rather than impaired diffusion, accounts for the lower measured substrate levels. This relationship is further illustrated by grouping the data according to droplet size (Fig. S11).

After 10 minutes, substrate concentration was consistently lower in active droplets than in enzyme-less droplets, while product concentration was higher across all size rages (Fig.S12), confirming that total substrate/product flux is faster in enzymatically active droplets.

To directly compare the initial stages of substrate diffusion before substantial substrate depletion occurred, we focused on droplets smaller than *<*40 *µ*m during the first 120 s following substrate addition (Fig. 3G). Throughout this period, active droplets accumulated substrate more rapidly than enzyme-free droplets. Because a fraction of the substrate was simultaneously converted into product, we also considered the combined concentration of substrate and product. Within the first 60 s, the total molecular concentration in active droplets was two- to three-fold higher than the substrate concentration measured in inactive droplets, demonstrating that substantially more substrate had entered the droplets. We concluded that the enhanced substrate diffusion is due to the activity-dependent reduction in apparent viscosity observed in Fig.2, as molecular diffusion and viscosity are inversely related through the Stokes–Einstein relationship. These results demonstrate that enzymatic activity not only remodels the physical properties of the crowded protein environment but also functionally enhances substrate diffusion, thereby mitigating diffusion limitations.

### Substrate-BSA interaction network underlies viscosity modulation

Having established that enzymatic activity reduces the apparent viscosity of crowded protein droplets and enhances substrate diffusion, we next sought to identify the molecular origin of this physical remodelling. We hypothesized that transient interactions between substrate molecules and the surrounding BSA network contribute to the apparent viscosity of the crowded phase, and that enzymatic turnover continuously remodels this interaction network. To explore this possibility, we first performed molecular docking analyses to characterize how each substrate interacts with the BSA-rich environment.

Following preparation of the BSA crystal structure, CavitOmix was used to identify potential surface-accessible binding cavities, 68 predicted cavities distributed across the protein surface (Table S2). Electrostatic surface mapping using APBS further revealed a heterogenous distribution of positively charged, negatively charged, and neutral regions (4A and Fig. S13), consistent with previous studies demonstrating that globular proteins present chemically diverse ligand-binding environments rather than a single dominant binding site.^58^ Together, the widespread distribution of predicted cavities and the heterogeneous electrostatic landscape indicate that BSA provides numerous spatially distinct interaction sites capable of accommodating small molecules through multiple weak, transient interactions. Such multivalent binding is expected to promote the formation of a dynamic substrate-protein interaction network.

The reaction products were then investigated for their differing ability to engage the distributed interaction sites identified on the BSA surface. Molecular docking was therefore performed for D-glucose, ONP, and DDAO against all 68 predicted binding cavities using AutoDock Vina, implemented through Docking Pie in PyMOL. Upto 20 binding poses were evaluated for each ligand-cavity pair. All three molecules displayed broad binding distributions across the protein surface, with D-glucose, ONP, and DDAO occupying 77%, 80%, and 84% of the predicted cavities, respectively (Fig. S14). The slightly broader binding profiles of ONP and DDAO are likely attributable for their aromatic moieties, which provide additional opportunities for hydrophobic and *ϱ*-mediated interactions with the chemically diverse BSA surface. Consistent with this, ONP and DDAO exhibited more favorable docking energies than D-glucose at the vast majority of binding sites (Fig. S14, Table S3), with only a small number of cavities deviating from this trend. The totaling of the binding energies across all accessible cavities fuether highlights the substantially greater interaction potential of the aromatic products, yielding cumulative binding free energies (ΔG_bind_) of −53.1, −177.1, and −191.2 kcal/mol for D-glucose, ONP, and DDAO, respectively. The corresponding average docking energies are presented in Fig. 4B. Together, these findings support a model in which the reaction products do not bind to a single high-affinity site, but instead engage in numerous weak, spatially distributed interactions across the BSA surface. Such transient multivalent interactions provide a molecular basis for the dynamic substrate-protein interaction network proposed to influence apparent viscosity of the crowded protein droplets, with these interactions being stronger and more abundant for ONP and DDAO than for D-glucose.

**Figure 4:**
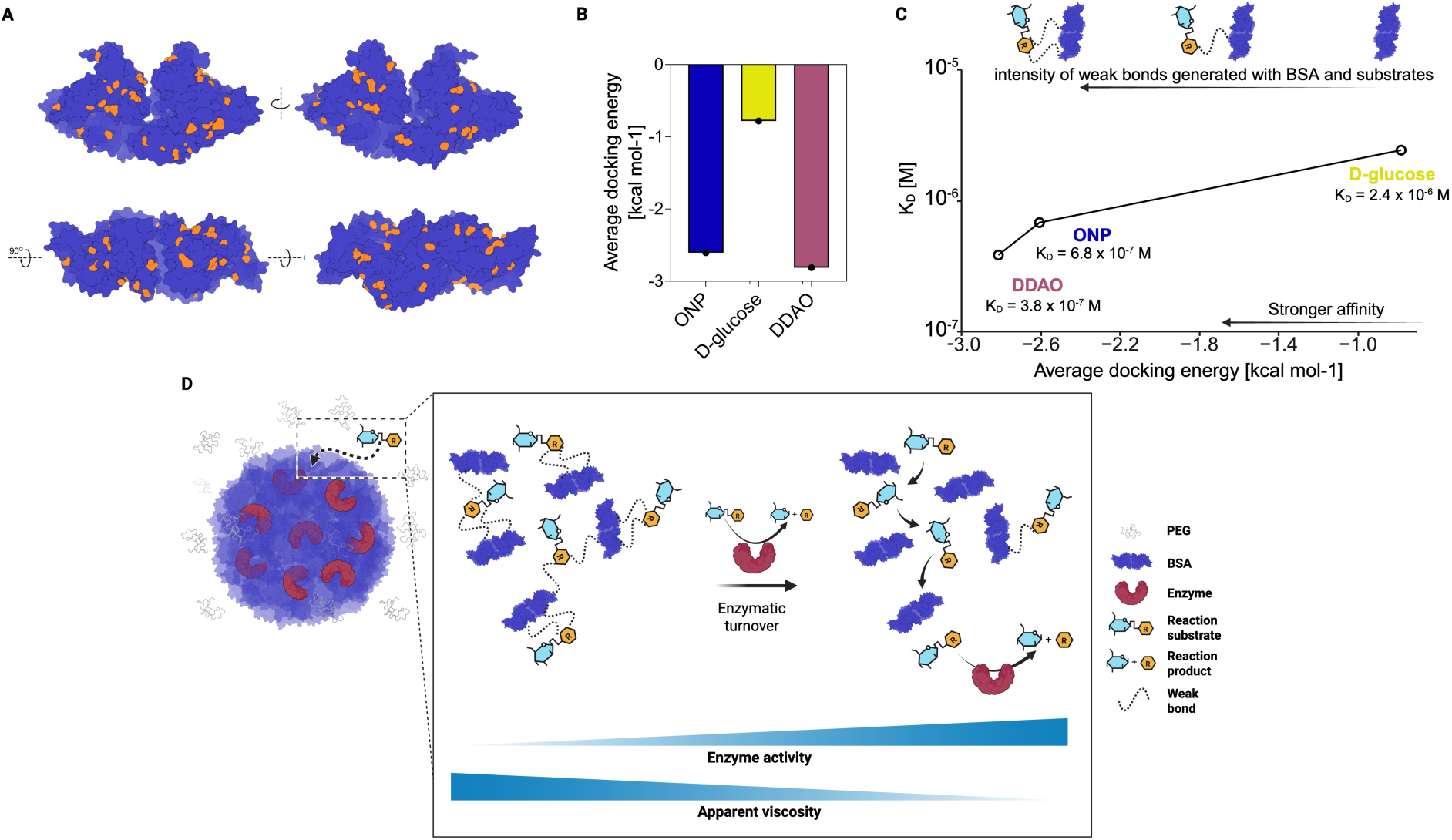
Product–BSA interactions build a network that affects the relative apparent viscosity. **A** Structural model of bovine serum albumin (BSA) showing the distribution of 68 potential binding cavities (highlighted in orange) across the protein surface, calculated using CavitOmix.^57^ **B** Average Docking energies (kcal/mol) of D-glucose, ONP, and DDAO over all 68 predicted BSA surface binding cavities, calculated using AutoDock Vina and visualized with Docking Pie. **C** Dissociation constant values (*K_D_*) of BSA from the SPR experiment support the molecular docking of the average BSA binding affinity for the different products (from panel B). **D** Working model illustrating how transient small-molecule–BSA interactions contribute to the apparent viscosity of crowded protein droplets and are perturbed during enzymatic turnover. This model is consistent with the observed reduction in apparent viscosity and enhanced substrate diffusion. This images was created with BioRender.com.

Surface Plasmon Resonance (SPR) analysis was used to experimentally validate these computational predictions, through the assessment of the binding affinity of the different reactions products to BSA. The interactions of five concentrations of the reaction products with BSA immobilised onto the SPR sensor surface are shown in (Fig.S15) and fitted with a multivalent (bi) model that closest represents multiple ligands (reaction products) binding to an analyte (BSA). The measure equilibrium dissociation constants (*K_D_*) closely followed the affinity ranking predicted by molecular docking, with DDAO exhibiting the strongest binding (*K_D_* = 3.81×10^−7^ M), followed by ONP (*K_D_* = 6.75×10^−7^ M), while D-glucose bound more weakly (*K_D_* = 2.42×10^−6^ M) (4C and Fig.S15). Thus, D-glucose displayed an approximately 3.6-fold lower affinity than ONP and a 6.4-fold lower affinity than DDAO. The close agreement between the docking predictions and the SPR measurements provides strongly with the BSA surface than D-glucose. Taken together, these independent computational and experimental analyses support the existence of a dynamic network of weak, multivalent substrate-protein interactions.

These complementary computational and experimental measurements support a model in which numerous weak substrate–BSA interactions collectively contribute to the apparent viscosity of crowded protein droplets (Fig. 4 D). During catalysis, substrate molecules are continuously converted into products, thereby dynamically remodelling the substrate-BSA interaction network and reducing the apparent viscosity of the crowded environment. This turnover-dependent remodeling provides a plausible molecular basis for the activity-dependent fluidization observed by microrheology and FRAP (Fig. 2) and the enhanced substrate diffusion observed in active droplets (Fig. 3).

Consistent with this mechanism, bulk shear rheology provided complementary evidence of activity-associated viscosity changes across the three reaction types, although interpretation is limited by the low droplet volume fraction and the macroscopic, steady-state nature of the measurement (see Supplementary Material and Methods and Fig. S16–S18).

## Discussion

This study demonstrates that enzymatic activity enhances substrate diffusion within crowded, phase-separated environments, with a reduction in apparent viscosity serving as the physical mechanism underlying this enhancement. Using *β*-galactosidase as a model enzyme and three chemically distinct substrates, we show that enzymatic turnover remodels the physical environment of the BSA-rich phase, thereby enhancing substrate diffusion up to three-fold under diffusion-limited conditions (diameter *<*40 um) compared to in enzyme-free droplets. Building on the PEG-BSA droplet platform we previously established, a recent parallel study using lactate dehydrogenase and alcohol dehydrogenase in the same system showed that enzymatic activity promotes fluidisation of the condensate, inferred from FRAP half-recovery times and nanoparticles diffusion coefficient.^38^ Our work complements and mechanistically extends those findings in three ways. First, we directly quantify the functional consequence for substrate availability, tracking real-time substrate and product diffusion inside droplets of defined size. Second, we generalised the phenomenon by using a novel enzyme and reaction type as well as detecting activity-dependent changes in the apparent viscosity of the crowded phase using particle-tracking microrheology and FRAP, with bulk rheology providing complementary evidence, grounding the diffusion enhancement in a measurable physical mechanism. Third, molecular docking and SPR affinity data pinpoint the molecular origin of the effect, showing that this viscosity modulation constitutes a positive feedback mechanism linking enzymatic activity to its own substrate supply.

We propose that enzymatic turnover enhances substrate diffusion by continuously remodelling transient non-covalent interactions between reacting small molecules and the protein crowder, thereby lowering local viscosity and easing molecular diffusion through the crowded phase. Molecular docking and SPR support this mechanism, showing that the tested substrates bind at multiple sites on the BSA surface with affinities that track the degree of viscosity reduction. These experiments are in agreement with the literature data since BSA is known to interact with several other aromatic molecules.^59–62^ Crucially, the FRAP data showed reduced T-halfs at increasing substrate (ONP-gal) concentrations that resemble the enzymatic turnover number, linking the amount of enzymatic activity to enhanced diffusion. This diffusion enhancement is most pronounced in small droplets (diameter *<*40 um), where the surface-to-volume ratio is highest and the diffusion constraints on the reaction are most severe; in larger droplets, enzymatic consumption of substrate before it can fully penetrate the droplet interior limits the benefit of reduced viscosity, and substrate availability instead becomes rate-limiting. In crowded environments like the cytosol, where the high concentration of protein-binding sites can otherwise slow molecular diffusion, enzymatic turnover may act as a diffusion regulator by continuously breaking weak substrate-protein bonds as the product is released, thereby reorganising the macromolecular network to keep substrate moving, an effect that should be most consequential in small, metabolically active compartments.

Diffusion of proteins and metabolites through the crowded cytosol is a fundamental determinant of cellular function, governing the availability of binding partners and the rate of molecular diffusion. For example, a bulk viscosity of only 5 mPa·s is sufficient to impede kinesin-1 movement along microtubules and thus its diffusion activity, with the magnitude of inhibition depending on crowder size.^63^ Because cellular viscosity appears constant across human cell lines and throughout the cell cycle,^64,65^ it is thought to be a homeostatically regulated parameter,^66,67^ and its dysregulation has been implicated in pathologial context as well, from cancer to neurogeneration.^68–70^ Our findings suggest that local, activity-dependent modulation of viscosity by enzymes offers cells a mechanism to tune diffusion and substrate availability locally, without altering bulk cellular viscosity; a route by which metabolic activity itself could regulate molecular diffusion in specific microenviroment. Although the PEG–BSA droplets used here provide a simplified model of the cytoplasm, they establish a physical principle by which metabolic activity could locally regulate molecular diffusion within a crowded protein environment.

The diffusion-enhancing mechanism identified here, substrate-crowder network disruption by catalytic turnover, is distinct from the chemical-gradient-driven hydrodynamic flows identified by Dindo et al., ^52^ which arise from phoretic interactions between enzyme-generated chemical gradients and confining surfaces and which also enhance diffusion within active condensate. These two mechanisms are not mutually exclusive: in living condensate, both substrate-crowder network disruption and gradient-driven flow may operate simultaneously and synergistically to enhance diffusion in the metabolically active environment.

Previous in vitro studies of enzyme-viscosity relationships have been limited to dilute buffer conditions^12,71^ where viscosity is too low to appreciably affect diffusion, or have reported changes in viscosity only related to the enzymatic degradation of the crowding agents themselves. ^72–74^ Our work is distinct from prior literature since the crowding agent (BSA) is not degraded during the reaction. Instead, enzymatic turnover changes the composition and interaction properties of the small-molecule population within an intact crowded protein phase, providing a molecular framework for understanding how enzymatic activity can reshape substrate diffusion and availability in crowded environments.

## Conclusion

In conclusion, this work expands the conceptual framework for enzyme function within the cell. Enzymes emerge not merely as passive catalysts governed by chemical kinetics, but as active regulators of their own substrate supply, capable of reshaping the diffusive properties of their environment to sustain catalytic turnover. By linking molecular-scale enzymatic turnover to a mesoscale reduction in apparent viscosity and, in turn to enhancement of substrate diffusion and availability, this study offers a new framework for understanding how metabolic flux is sustained under the diffusion constraints imposed by macromolecular crowding. This work points toward a broader principle: metabolically active condensates are not passive crowded environments, but dynamic, self-regulating systems in which catalytic activity itself continuously reshapes the physical landscape that governs substrate availability.

## Materials and methods

### Materials

Polyethylene glycol (PEG) 4000 Da (A16151) was purchased from Alpha-Aesar. Bovine serum albumin (BSA) (A7638), 3-(Trimethoxysilyl)propylmethacrylate (440159), 2-Hydroxy-4’- (2-hydroxy ethoxy)-2-methylpropio phenone (410896), Poly (ethylene glycol) diacrylate (PEGDA) 700 Da (455008), PEGDA 10 kDa (729094), D-glucose (G8270), *β*-galactosidase (G5635), potassium phospate dibasic trihydrate (60349), horse-radish peroxidase (P8375), D-lactose (61339), dipotassium phosphate, disodium phosphate, ethanolamine, 1-ethyl-3- (3-dimethylaminopropyl)carbodiimide (EDC), hydrocloric acid, glycine, potassium chloride, sodium acetate, sodium chloride, and Tween20 were purchased from Sigma-Aldrich. Alexa Fluor 488 Microscale Protein Labeling Kit (A30006), Fluoro-Max red fluorescent particle (R0100), (9H-(1,3-Dichloro-9,9-Dimethylacridin-2-One-7-yl) *β*-D-Galactopyranoside) (D6488) were purchased from Thermo Fischer Scientific. 9H-(1,3-dichloro-9,9-dimethyl acridin-2-one) (DDAO) was purchased from Setareh Biotech. Potassium phosphate monobasic (42420) was purchased from Acros Organics. Deuterium oxide (D2O) (DE50B) was purchased from Apollo. N,N-dimethylformamide (DMF) (D119) was purchased from Fisher Chemicals. O-dianisidine (119-90-4) was purchased from Tokyo Chemical Industry. ONP-gal (25027-71), sodium hydroxide (31511-05), magnesium chloride (20909-55) and tris (hydroxymethyl) aminomethane ( TRIS, 35406-75), Dimethyl Sulfoxide (DMSO, 08904-85), o-nitrophenol (ONP, 24919-92), glucose oxidase (16831-14) were purchased from Nacalai Tesque. Potassium chloride (163-03545) was purchased from Fujifilm Wako.

### PEG-BSA system optimization and characterization

The droplet solution was optimized in the following buffer conditions: Tris (hydroxymethyl) amino-methane-hydrochloride (TRIS-HCl) 300mM buffer pH 7.0, in the presence of potassium chloride 50 mM, Magnesium dichloride 2 mM, BSA 447 *µ*M and 18% PEG (w/w). Importantly, PEG solution was added last and gently mixed to obtain the BSA droplets solution. The system was characterized by measuring the droplet volume fraction and the concentration of PEG and BSA in both the droplet and PEG-rich phase. In order to measure the droplet volume fraction, 1 ml of droplet solution was prepared and centrifuged for 60 mins and 16000 rcf. The PEG-rich phase (supernatant) was removed and the remaining droplets (bottom phase) were centrifuged for 20 mins at 16000 rcf. Remove the residual PEG-rich phase from the centrifuged droplets. 50 *µ*l of water were added to the droplets and mixed by pipetting. This process dissolves the dense droplets phase and allows the measurement of the volume by pipetting. The BSA and PEG concentrations in the two-liquid phases were quantified using UV-Vis spectrophotometry and NMR as previously reported.^40^

### Protein Surface Binding Cavity Analysis

Potential binding cavities across the surface of bovine serum albumin protein (BSA, PDB code: 4F5S) were calculated using CavitOmix (v1.0, 2022, Innophore GmbH). The corresponding hydrophobicity module of the program VASco^57^ was used to analyze the hydrophobicity of the cavities. The cavities were calculated using a modified LIGSITE algorithm.^75^ The settings include a minimum of five site points and a maximum of 100 site points per site. The sites generated were produced with a grid spacing of 0.7 between potential binding cavities, with a total of 68 calculated potential binding cavities identified. The surface of the protein was analyzed by using APBS (v3.4.1), with the protein initially prepared using PDB2PQR (v3.6.1) at a pKa of 7.0 and an AMBER forcefield, ensuring that waters are removed, new atoms not rebuilt too close to current atoms and optimizing the hydrogen bonding network.^76^ Adaptive Poisson-Boltzmann Solver (APBS) was then subsequently used with mg-auto calculation time, to determine electrostatic surface visualization.^76^

### Docking Calculations

For each of the calculated potential surface binding cavity, docking calculations on the three ligands (D-glucose, ONP and DDAO), were performed using AutoDoc Vina (v1.2),^77,78^ via Docking Pie (v1.2)^79^ a Molecular Docking plugin for PyMOL (The PyMOL Molecular Graphics System, Version 3.0 Schrodinger, LLC.). The receptor was generated with the addition of hydrogens and removal of water molecules, while the ligand was generated with all active torsions. Grids settings were determined using the calculated potential binding cavities, with Grid Center Coordinates and Grid Dimensions shown in Supplementary Information (Table S2). Docking was performed with an exhaustiveness of 8 and an energy range of 3. Up to 20 poses per input ligand structure were saved for each docking run, to generate a large number of diverse poses.

### Binding and Affinity Performace by Surface Plasmon Resonance (SPR)

Immobilization of the BSA protein onto the SPR sensor surface occurs through a PEG coated Au chip that was initally pre-conditioned by a PBS pH 7.4 and 0.01% Tween 20 running buffer at 10 *µ*L min^−1^ within the SPR. A 1 mL aqueous solution containing 40 mg EDC and 10 mg NHS was passed over the chip ( 6 min at 10 *µ*L min^−1^). 40 *µ*g of BSA in 1 mL of running buffer and 10 mM of sodium acetate were injected over the left channel (working channel) of the chip for 1 min. An 8 min injection over bother channels (working and reference) of quenching solution (1 M ethanolamine, pH 8.5) was added, followed by a continuous flow of running buffer at 10 *µ*L min^−1^. all injections were taken from a stable baseline.

Kinetic analysis in binding of the analytes (reactions products) to BSA were performed using an SPR assay coinsisting of 2 min association phase (PBS and 0.01% Tween20 containing analyte at concentrations of 3.125-50 nM), followed by a 5 min dissociation phase (running buffer only). After each binindg cycle, the sensor surface was regenerated using 10 mM glycine-HCl (pH 2) for 1 min and subsequently stabilised with running buffer for a further 1 min. A blank injection of running buffer was performned prior to analyte measurements, with analyte concentrations introduced dequentially in increasing order.

Following data acquisition, signals from the reference channel were subtracted from those of the working channel to correct for non-specific responses. All rebinindg measurements were performed in triplicate. Sensorgrams were analysed using the Biacore T200 Evaluation Software and fitted to a Langmuir bivalent binding model. The association rate constant (k_a_), dissociation rate constant ((k_d_), and maximum binding capacity (B_max_) were determined by global fitting, while the response signals were fitted locally. Equilibrium dissociation constants (K_D_) were calculated from the ratio k_d_/(k_a_.

### Particle-tracking microrheology

During the droplet preparation as described before, red fluorescent beads of 1.1 *µ* m diameter (Fluoro-Max R0100,Thermo Fisher, USA) were incorporated into the droplet preparation mixture at a final dilution of 1/1000 prior to the addition of PEG. This process ensures the selective localization of the fluorescent beads into the droplet phase. The droplets solution were diluted 1/10 and 20 *µ*L of droplets were placed in a sample chamber^52^ to be imaged after the addition of 180 *µ*L of PEG-phase containing the selected amount of reaction substrate. The particles were visualized using a confocal microscope (Zeiss LSM880 Airyscan) with a 63X objective and the particles were excited using a laser of *λ_g_* = 561 nm. The images were recorded on a region of interest (ROI) of 512 x 512 pixels. Multiple fields were imaged to ensure a total of at least 50 particles per condition. The images were captured at at 25*^°^*C, under a frame rate of 2.12 frames per second for up to 200 s..

Particle trajectories were analyzed using the TrackMate plugin to extract coordinate traces, and mean squared displacement (MSD) curves were computed with TrackMateR as:

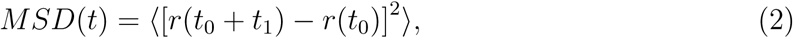

where *r* is the particle position and angle brackets denote averaging over all particles. Diffusion coefficients (*D*) were estimated from the slope of the MSD, assuming a free 2-D diffusion:

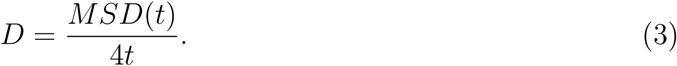

Viscosity (*ϑ*) was then calculated using the Stokes–Einstein– Sutherland relation:

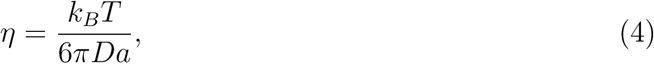

where *k_B_* is Boltzmann’s constant, *T* is temperature in Kelvin, and *a* is the particle radius.

### MathLab script

Analysis of droplet fluorescence was conducted using self-made Matlab scripts. The elliptical circumference of unmoving droplets was extrapolated from a user-defined selection of points. Then, the concentrations of product and substrate were calculated using the experimentally determined calibration curve. For average concentration data, only pixels inside the circumference were considered. The scripts used in the analysis are freely available upon request.

## Supporting information

Supplementary Information file including supplementary images and tables

## Acknowledgement

The research was supported by the Okinawa Institute of Science and Technology Graduate University (OIST) with subsidy funding to P.L., and A.Q.S. from the Cabinet Office, Government of Japan. P.L. thanks the financial support from Takeda Grant. P.L. thanks METX JSPS Kakenhi 24K01995. M.D. thanks the financial support from Japan Society for the Promotion of Science (JSPS) for the Kakenhi Early Career Scientist N.22K15065. The authors thank the fruitful discussions with Dr. Vincenzo Calabrese, Ms. Erika Fukuhara, Dr. Yusran Abdillah Muthahari, and Ms. Arisa Yokokoji. The authors thank Prof. Marco Edoardo Rosti for the useful comments on the data and for the critical reading of the text. We are grateful for the help provided by the imaging section at OIST; in particular, we thank Paolo Barzaghi for the support with the confocal microscopes. Declaration of Generative AI and AI-assisted Technologies: In preparing this manuscript, the authors used ChatGPT (OpenAI) to assist with scripting for data plotting and to identify grammatical mistakes in the text. The suggestions were manually reviewed and selectively incorporated into the text. The authors take full responsibility for the content of the publication. A.B. and M.R. would like to thank A. Toriyama for his works that inspire us to keep pushing forward. The images of the present work were prepared using Biorender.com.

## Supporting Information Available

The supporting information contains the experimental procedures, reaction schemes, and supporting data of this study. Experimental movies are available as web-enhanced objects.

