## Supplementary Information file including supplementary images and tables for "Enzyme Activity Regulates Substrate Diffusion by Modulating Viscosity in Crowded Milieu"

### SUPPORTING INFORMATION for Enzyme Activity Regulates Substrate Diffusion by Modulating Viscosity in Crowded Milieu

**2 Supplementary results 12**

**3 Supplementary movies 35**

### Experimental section

#### Enzyme partitioning inside the droplets

To evaluate the partitioning of $\beta$-galactosidase in the PEG-rich phase and in the droplet phase, we measured the enzyme activity in both liquid phases. Specifically, 0.3 mL of droplet mix containing 20 nM $\beta$-galactosidase (0.6 $\mu$M inside the droplets) was centrifuged for $\geq$ 30 minutes at 16000 g at room temperature. Then, the supernatant (the PEG-rich phase) was transferred to a new tube while the bottom phase (the droplets) was subjected to another centrifugation cycle. The supernatant was removed again from the centrifuged droplets. The droplets and supernatant solutions were diluted 1:2 using TRIS-HCl buffer (TRIS-HCl 0.3 M, pH 7.0 containing 2 mM magnesium dichloride and 50 mM potassium chloride). Then, reaction timepoints were taken by adding 3 $\mu$L of droplets/PEG-phase into 147 $\mu$L of TRIS-HCl buffer containing 5 mM ONP-gal; the reaction was stopped with heat (100 C for 45 seconds) and then centrifuged at 13000 g for 3 minutes. 100 $\mu$L of the samples were moved into a 96-well plate and the absorbance at 420 nm was recorded using Multiskan Go plate reader (Thermo Fisher Scientific). The absorbance was converted into product (ONP) concentration by using the Beer-Lambert law using 4500 $M^{-1}cm^{-1}$ as ONP molar attenuation coefficient ($\varepsilon$) and 0.29 cm as optical path-length ($l$). Moreover, $\beta$-galactosidase partitioning inside the droplets was assessed using confocal microscopy to confirm the absence of protein aggregation. The enzyme was labeled using the protein labeling kit Alexafluor-488 and observed using a Spinning Disk Confocal (Nikon, Andor CSU) microscope with 63$\times$ oil immersion lens. Specifically, droplet solution containing 0.5 $\mu$M 488-labeled $\beta$-galactosidase in the mix ($\approx$ 15$\mu$M inside the droplets) were diluted 1:100 with the PEG-rich phase and imaged on top of a (700 Dalton) PEGDA-coated glass slides[1].

#### Enzyme kinetics assays

The kinetic parameters (Kcat and Km values) of $\beta$-galactosidase were assessed in the PEG-rich phase and in the droplets + PEG-rich phase. The activity of $\beta$-galactosidase was assayed on three substrates: 4-nitrophenyl $\beta$-D-galactopyranoside (0.35-20 mM), D-lactose (0.5-50 mM) and 9H-1,3-Dichloro-9,9-Dimethylacridin-2-One-7-yl $\beta$-D-galactopyranoside (0.25-3.5 mM). Droplets and PEG-rich solution containing 100-200 nM enzyme were prepared for the kinetic test. Note that inside the droplets the local concentration of enzyme increases 30 times since the droplets represent the 3.3% of the volume fraction (Fig.S2). Thus, the final local concentration of the enzyme inside the droplet is 3-6 $\mu$M. The solutions were diluted (with the PEG-rich phase + substrate) at the final enzyme concentration of 0.25-1 nM, while the local enzyme concentration inside the droplets remains 3-6 $\mu$M. Diluting the enzyme-droplets solution makes the droplets smaller and prevents the diffusion of the substrate becoming the rate-limiting step of the enzymatic reaction (Fig. S8). At each time point, the reaction mix was stopped using heat and stored on ice protected from light. The samples were then centrifuged and the amount of product in the reaction mix was determined by measuring the products' absorbance using a Multiskan Go plate reader (Thermo Fisher Scientific) and a calibration curve.

The $\beta$-galactosidase kinetics values on the substrate ONP-gal (1M stock in 100% DMSO, stored at -20°C) were determined by using a droplets mix containing 200nM enzyme (6 $\mu$M inside the droplets) diluted 1:700 using the PEG-phase for a final enzyme concentration of 0.286 nM in a total reaction volume of 1500 $\mu$L. The reaction time-points were taken by removing 300 $\mu$L of the reaction mix and heating them for 60 seconds at 100 °C, and stored on ice. The time-point tubes were centrifuged (16000 g for 3 minutes) and 200 $\mu$L of the reaction mix were placed in a spectrophotometer to determine the product (ONP) absorbance at 420 nm (Fig.S4).

The kinetics values on the substrate D-lactose (0.6 M D-lactose in water, stored at -

20°C) were assessed by using the coupled assay method using glucose oxidase, horse radish peroxidase, and o-dianisidine.[2–4] In details, a droplets mix containing 100nM enzyme (3 $\mu$M inside the droplets) diluted 1:100 using the PEG-phase for a final enzyme concentration of 1 nM in a total reaction volume of 900 $\mu$L. The reaction time-points were taken by removing 100 $\mu$L of the reaction mix and heating them for 45 seconds at 100°C, and stored on ice. The time-point tubes were centrifuged at 16000 g for 3 minutes. Then, 20 $\mu$L of the reaction mix were added to 180 $\mu$L of detection mix (200 $\mu$M o-dianisidine, 1 $\mu$M D-glucose oxidase, 1 $\mu$M horse radish peroxidase in potassium phosphate buffer 0.1M, pH 7.0), and incubated for 20 mins at room temperature, protected from light. Then, 200 $\mu$L of the solution was placed on a spectrophotometer to determine the absorbance of the product of the coupled assay (oxidased o-dianisidine) at 450 nm (Fig.S5).

The $\beta$-galactosidase kinetics values on the substrate DDAO-gal (200 mM stock in 100% DMSO, stored at -20 ° C) were determined using a droplet mix containing 100 nM enzyme (3 $\mu$ M within the droplets) diluted 1:400 using the PEG-phase (+ substrate) for a final enzyme concentration of 0.25 nM in a total reaction volume of 125 $\mu$L. The reaction time-points were taken by removing 25 $\mu$L of the reaction mix and heating them for 45 seconds at 100 °C, and stored on ice. The time-point tubes were centrifuged (16000 g for 3 minutes) and 10 $\mu$L of the reaction mix were placed 190 $\mu$L of TRIS-HCl buffer 0.3M (pH 7.0) to determine the product (DDAO) absorbance at 648 nm (Fig. S6). The substrate DDAO-gal showed precipitation in concentrations greater or equal than 1 mM, as tested in the kinetic assay. Thus, the precipitated fraction was accounted and the data of the Michaelis-Menten were normalized accordingly (Fig. S20). Specifically, the precipitated fraction was determined by using a calibration curve of DDAO-gal concentrations which do not show precipitation (0-0.5 mM). The standard solutions were prepared in the PEG-rich phase. Then, 5 $\mu$L of the standard solution were moved into a cuvette containing 245 $\mu$L of TRIS-HCl 0.3 M (pH 7.0) and the samples' absorbance was recorded at 464 nm using a UV-1900 (Shimadzu) spectrophotometer (Fig. S20A). Then, the DDAO-gal samples were prepared at concentrations that show precipitation (1, 2, 5, and 10 mM) in the PEG-rich phase. The samples were centrifuged for 5 minutes at 16000 g to remove the precipitated DDAO-gal. Then, 5 $\mu$L of the supernatant was added to a cuvette containing 245 $\mu$ L of 0.3 M TRIS-HCl (pH 7.0) and the absorbance was recorded at 464 nm (Fig. S20B).

The reaction products ONP (50 mM stock in 50% DMSO and 50% TRIS-HCl 0.3 M pH 7.0, stored at +4°C), D-glucose (500 mM stock in water, stored at 25 °C), and DDAO (100 mM in 100% DMSO, stored at -20 °C) were used to make the calibration curve in the PEG-rich phase to convert the absorbance value into product concentration (Fig. S4B, S5B and S6B). These data were analyzed using GraphPad Prism and fitted using the Michaelis-Menten equation.[5]

#### FRAP experiments

FRAP experiments were conducted using a Nikon A1R system on a Nikon TiE2 inverted microscope using a 100x oil immersion objective (100x/1.49 Oil Apo TIRF). The droplets used for this experiment were prepared containing a 15% of BSA fraction labeled with the Alexafluore 488 protein labeling kit (67 $\mu$M of 488-labeled BSA and 380 $\mu$M BSA), as previously reported.[1] The enzyme concentration range tested in the FRAP experiment was 0-0.1 $\mu$M in the system (and 0-3 $\mu$M inside the droplets). Then, 2 $\mu$L of droplet mix was diluted into 18 $\mu$L of PEG phase, and added to the sample chamber.[1] Subsequently, 180 $\mu$L of PEG-rich phase containing 25 mM ONPG were added to the droplets mix and the solution was incubated for 5 minutes to allow the substrate to diffuse inside the droplets and activate the enzyme. The droplets were imaged for 120 seconds (220 frame total) using a $\lambda_g$= 488 nm laser line by adjusting the laser power (0.2-0.5%) to obtain a droplets average fluorescence intensity of 1500-3000 A.U. The droplets were first imaged for 10 seconds to record the "pre-bleach" images, followed by 1 bleach frame with the laser power locally intensified to 0.5-1.5% to obtain a bleach depth intensity of ≈80%, and finally 200 post-bleach frames with the initial laser power intensity. For this analysis, only droplets of a diameter of 10-35 $\mu$m were used to avoid any possible substrate diffusion limitations. The videos were recorded after maximum 10 minutes from the addition of the reaction substrate, then a new droplets sample was set up. The movies were analyzed using Fiji ImageJ[6] to determine the average intensity over time of the bleached region of interest (ROI), a reference ROI (consisting of a droplets area not subjected to bleaching), and a background ROI (an area outside of the droplets). The ROI analysis was then exported and processed with EasyFRAP online tool[7] using double normalization method and "double" as curve fitting equation parameter; if an error was displayed in the curve fitting, the latter parameter was changed to "single". Only sample with a mobile fraction higher than 40% and with fitting curve with $R^2 \geq 0.85$ were considered for the analysis and plotted in Fig.3D.

#### Calibration curve of DDAO-Gal and DDAO at confocal microscope

500 $\mu$L aliquots of BSA droplets solution were prepared and a range of substrate/product concentration (0-200 $\mu$M) were added. The droplets were left two hours at room temperature to assure to reach diffusion equilibrium in the droplets/PEG phase. Then, the droplets pelleted on the bottom of the tube were resuspended with the pipette tip. 2 $\mu$L of droplets were added to 18 $\mu$L of PEG phase. The diluted droplets were moved into a sample chamber placed on a coated glass;[1] after adding 180 $\mu$L of PEG phase, the dataset was recorded with the confocal microscope LSM880 (20x lens) by exciting the probes at $\lambda_g$ = 488 nm and $\lambda_g$ = 646 nm and by recording the signal at the range of $\lambda_g$ = 588-624 nm and $\lambda_g$ = 657-680 nm for DDAO-gal and DDAO, respectively. The signal was stable for several minutes before decreasing due to bleaching/diffusion outside of the droplets (data not shown). The droplets intensity at different substrate/product concentration was measured using ImageJ software and plotted to build a calibration curve using Graphpad Prism (Fig. S9).

#### Diffusion experiment of DDAO and DDAO-gal in droplets

Droplets without enzyme were prepared and centrifuged for 20 seconds at 10000 rcf and then resuspended with a pipette tip to obtain bigger droplets. Resuspended droplets (2 $\mu$L) were diluted 1:10 with PEG phase, moved into a sample chamber placed on coated glass[1] and imaged by using Zeiss LSM880 microscope. Later, 180 $\mu$L of PEG phase containing 0.22 mM DDAO-galactoside or 0.22 mM DDAO were added to the diluted droplets (0.2 mM final concentration). Then, the diffusion of substrate and product inside the droplets of different size have been recorded for 1400 seconds using 488-nm, 648-nm laser and bright field channels (Supporting movies 9). The same experiment was repeated by using droplets containing 9 $\mu$M $\beta$-galactosidase and by adding PEG phase solution containing 0.22 mM DDAO-Gal (0.2 mM final concentration) during the imaging process. The fluorescence intensity was converted into product/substrate concentration by using a calibration curve (Fig. S9). Different not-linear diffusion patterns were shown (Fig.S19). We hypothesized that the patterns are due to the different focus planes and/or because droplets are not spherical but pinned on the bottom[1] which can distort concentration profiles and result in non-linear diffusion. For these reasons, only the experiments showing linear diffusion patterns reaching the product/substrate concentration added to the solution were considered for the analysis. depending of the sample group, 0-50% of the tracks were discarded due to non-linearity.

#### Bulk rheology experiments

To evaluate how enzymatic activity influences the macroscopic rheology of a crowded biphasic system, we measured the apparent shear viscosity of the PEG-BSA mixture using shear rheometry. This *bulk solution* comprises a PEG-rich continuous phase (which contains one of three reaction substrates: ONP-gal, lactose, or DDAO-gal) and dispersed BSA-rich droplets (reaction scheme in Table S1). In this set-up, the enzyme $\beta$-galactosidase is localized within the BSA-rich droplets, while the substrate is added in the PEG-rich phase and rapidly diffusing in the BSA-rich droplets phase.

We define the *relative viscosity* $\eta^*$ as:

$$\eta^* = \frac{\eta}{\eta_0}, \tag{S1}$$

where the apparent shear viscosity $\eta$ under each condition is normalized by the baseline viscosity $\eta_0$, defined as the viscosity of the control system (PEG-rich continuous phase with substrate and BSA-rich droplets lacking enzyme). Both $\eta$ and $\eta_0$ were measured under identical shear conditions ($\dot{\gamma} = 1$ s$^{-1}$, $T = 24$°C). This normalization isolates the effect of enzymatic catalysis from changes due to solvent composition or droplet volume fraction.

Steady flow shear rheological measurements were conducted using a strain-controlled rheometer (ARES G2, TA Instruments, USA). The shear viscosity was measured at T = 24°C over a shear rate range of $0.1 < \dot{\gamma} < 20$ s$^{-1}$using a 40 mm (in diameter) stainless-steel cone-plate geometrywith 1 degree angle which minimizes confinement effects and ensures uniform shear distribution. Prior to measurement, the bulk solution was diluted by a factor of 1/10 in PEG phase to maintain consistency across samples and control for droplet volume fraction (Fig.S16).

We next assessed how enzymatic activity alters the shear viscosity of the bulk solution. BSA droplets in the presence (active enzyme condition) or absence of 10 nM $\beta$-galactosidase in the bulk (equivalent to 300 nM in-droplet concentration) were introduced into PEG solutions containing either substrate (25 mM ONP-gal, 20 mM D-lactose, or 1 mM DDAO-gal) or the corresponding reaction product (25 mM ONP, 20 mM D-glucose, or 1 mM DDAO) (Fig.S17A).

As shown in Figure S17A enzyme-containing systems exhibited a marked reduction in $\eta$ across all substrates in droplets+continuous phase: from 0.0316 ± 0.0042 to 0.0153 ± 0.0048 Pa·s for ONP-gal, from 0.0469 ± 0.0054 to 0.0279 ± 0.0101 Pa·s for D-lactose, and from 0.0432 ± 0.0079 to 0.0142 ± 0.0053 Pa·s for DDAO-gal. These reductions indicate a strong coupling between enzymatic activity and viscosity reduction, with the largest effects observed for substrates that are more hydrophobic or have higher affinity for BSA. Products in the PEG-rich solvent with BSA droplets without enzyme showed an increase in relative viscosity compared to the lower values observed after enzyme addition, particularly for ONP and DDAO (Fig. S17A). Importantly, the relative viscosity ($\eta^*$) changes, defined as the bulk viscosity values of the droplets+continuous phase with enzyme activity normalised by the bulk viscosity value of the corresponding system with substrate but without enzyme, show a decrease that relates to the binding affinity (Fig.S17B) from the docking analysis, resembling the trend of the SPR data of Fig.4. These observations suggest that product–BSA interactions contribute to viscosity modulation. Our findings indicate that this enzymatic–viscosity coupling reorganizes macromolecular crowding within the droplets.

To distinguish the effect of enzymatic activity from the mere presence of the enzyme, we first examined enzyme-only controls (no substrate) across a range of concentrations (0, 0.1, 1, and 10 nM in bulk; 30–300 nM within BSA droplets). As shown in the inset of Fig.S19, relative viscosity ($\eta^*$) remained close to baseline (near 1.0) across all concentrations, with a slight decrease to $0.75 \pm 0.11$ only at the highest enzyme level (10 nM). These results confirm that enzyme alone has little impact, especially at low concentrations, on the bulk viscosity of the PEG solvent–BSA droplet system. When the ONP-gal substrate was included in the system, a clear enzyme activity-dependent reduction in $\eta^*$ was observed (Fig.S18). These results indicate that enzymatic catalysis, rather than the mere presence of the enzyme, is responsible for the observed changes in viscosity. However, the droplet system in the presence of enzymatic activity show some challenges for the bulk rheology approach since (i) the bulk rheology is a measurement of a stady-state system and not a dynamic system were the enzyme is biochemically active, (ii) the majority of the bulk solution is composed by PEG-phase (only about 3-0.03% is composed by droplets), (iii) the amount of droplets affects the viscosity (Fig.S16), (iv) the reaction is localised only in the droplets which can generate other effects by not distributing the reaction in the bulk, (v) droplets sedimentation and merging can affect the measured bulk viscosity. Due to these challenges, the bulk rheology measurements of the droplet system (PEG-continuous phase + 0.3% droplets) are less informative but still resemble the general trend of decreased viscosity in the presence of enzymatic activity, compared to the control samples with only substrate, product or enzyme (Fig.S16-18), in accordance to the previous results.

#### Steady flow shear rheology measurements

Steady-state shear rheology measurements were performed using a strain-controlled rheometer (ARES G2, TA Instruments, USA). Flow curves were acquired at 24°C over a shear rate range of $0.1 < \dot{\gamma} < 20$ s$^{-1}$ using a cone-and-plate geometry (40 mm diameter, 1 angle) to minimize confinement effects and ensure uniform shear distribution.

In the first set of experiments, we evaluated the impact of enzymatic activity in the presence of different substrates. Shear viscosity was measured for the full bulk system (PEG-rich phase + BSA-rich droplets) in the presence and absence of 10 nM $\beta$-galactosidase (corresponding to 300 nM enzyme within the droplets). Droplets were diluted by a dilution factor of 1/10 in PEG-rich phase, yielding a final enzyme concentration of 1 nM in the bulk. Substrate-saturated solutions were prepared with 25 mM ONP-gal, 20 mM D-lactose, or 1 mM DDAO-gal. For comparison, measurements were also conducted with corresponding reaction products (25 mM ONP, 20 mM D-glucose, and 1 mM DDAO). In a second set of experiments, we investigated the effect of varying enzyme concentrations. The droplet volume fraction was held constant while adjusting the enzyme concentration in the bulk to 0, 1, 10, or 100 nM, corresponding to 0, 0.03, 0.3, and 3 $\mu$M enzyme inside the droplets, respectively. Prior to each measurement that involves active enzyme, samples were incubated for 10 minutes to allow the enzymatic reaction to reach steady state and to minimize diffusion limitations.

### Supplementary results

Table S1: Overview of the different enzyme reactions tested in this study in the PEG-BSA droplet system.

| Reaction substrates added | Enzyme | Reaction products |
|---|---|---|
| ONP-gal | $\beta$-galactosidase | ONP + D-galactose |
| D-lactose | $\beta$-galactosidase | D-glucose + D-galactose |
| DDAO-gal | $\beta$-galactosidase | DDAO + D-galactose |

Table S2: Potential binding cavity grid coordinate centers in $\mathring{A}$, defined by inner box size in corresponding parentheses.

| Binding Cavity Sites | Grid Coordinates | | |
|---|---|---|---|
| | x | y | z |
| 1 | 41.97 (4) | 27.54 (4) | 63.72 (4) |
| 2 | 82.77 (4) | 35.74 (4) | 96 (1) |
| 3 | 57.02 (1) | 8.94 (4) | 92.42 (4) |
| 4 | 15.01 (4) | 39.21(1) | 105.18 (4) |
| 5 | 70.55 (4) | 26.04 (4) | 90.71 (1) |
| 6 | 65.24 (4) | 49.03 (4) | 103.04 (4) |
| 7 | 58.99 (1) | 27.61 (4) | 66.53 (4) |
| 8 | 57.98 (4) | 12.79 (1) | 74 (4) |
| 9 | 81.17 (4) | 20.11 (4) | 100.9 (1) |
| 10 | 26.12 (4) | 47.77 (4) | 94.1 (4) |
| 11 | 54.01 (1) | 6.85 (4) | 78.77 (4) |
| 12 | 25.08 (4) | 30.19 (1) | 82.37 (4) |
| 13 | -3.31 (4) | 28.05 (4) | 126.84 (1) |
| 14 | 39.3 (4) | 15.17 (4) | 69.48 (4) |
| 15 | 13.5 (1) | 9.25 (4) | 128.68 (4) |
| 16 | 17.53 (4) | -2.1 (1) | 126.86 (4) |
| 17 | 44.02 (4) | 7.03 (4) | 116.56 (1) |
| 18 | -0.78 (4) | 15.34 (4) | 95.96 (4) |
| 19 | 92.7 (1) | 18.02 (4) | 83.03 (4) |
| 20 | 61.98 (4) | 7.89 (1) | 72.61 (4) |
| 21 | 66.53 (4) | 47.31 (4) | 113.88 (1) |
| 22 | 67.57 (4) | 17.6 (4) | 69 (4) |
| 23 | -23.07 (1) | 28.97 (4) | 116.03 (4) |
| 24 | 4.43 (4) | 27 (4) | 131.27 (4) |
| 25 | 67.75 (4) | 3.28 (1) | 87.46 (4) |
| 26 | -1.44 (1) | 19.64 (4) | 83.12 (4) |
| 27 | 60.57 (4) | 48.12 (1) | 121.21 (4) |
| 28 | 78.59 (4) | 33.13 (4) | 104.3 (1) |
| 29 | 4.74 (1) | 9.76 (4) | 106.63 (4) |
| 30 | 40.19 (4) | 3.42 (1) | 119.49 (4) |
| 31 | -5.6 (4) | 29.85 (4) | 75.44 (1) |
| 32 | 40.65 (4) | 44.38 (4) | 119.23 (4) |
| 33 | 2.71 (1) | 4.95 (4) | 122.13 (4) |
| 34 | 6.39 (4) | 41.72 (1) | 103.13 (4) |
| 35 | 89.09 (4) | 24.87 (4) | 75.23 (1) |
| 36 | 45.35 (4) | 37.3 (4) | 127.1 (4) |
| 37 | 11.2 (1) | -1.7 (4) | 119.75 (4) |
| 38 | 5.92 (4) | 4.09 (1) | 106.62 (4) |
| 39 | 46.61 (4) | 6.59 (4) | 82.66 (1) |
| 40 | 0.64 (4) | 14.51 (1) | 126.57 (4) |
| 41 | 70.51 (4) | 27.24 (4) | 72.29 (1) |
| 42 | 75.98 (4) | 41.94 (4) | 96.03 (4) |
| 43 | -3.95 (1) | 14.64 (4) | 123.45 (4) |
| 44 | -10.78 (4) | 21.56 (1) | 96.67 (4) |
| 45 | 15.4 (4) | 33.81 (4) | 122.57 (1) |
| 46 | 62.77 (4) | 30.02 (4) | 70.86 (4) |
| 47 | 50.71 (1) | 44.8 (4) | 119.16 (4) |
| 48 | 2.72 (4) | 12.91 (1) | 95 (4) |
| 49 | 41.84 (1) | 23.8 (4) | 122.97 (4) |
| 50 | -0.17 (1) | 43.84 (4) | 107.54 (4) |
| 51 | 8.4 (4) | 41.56 (1) | 94.06 (4) |
| 52 | 11.64 (4) | 38.76 (4) | 89.25 (1) |
| 53 | 83.39 (4) | 37.19 (4) | 79.8 (4) |
| 54 | 61.95 (1) | 10.5 (4) | 68.69 (4) |
| 55 | 72.36 (4) | 30.97 (1) | 94.94 (4) |
| 56 | 18.7 (4) | 44 (4) | 92.2 (1) |
| 57 | -10.1 (4) | 32.7 (4) | 93.3 (4) |
| 58 | 42.12 (1) | 10.97 (4) | 128.33 (4) |
| 59 | 10.38 (4) | 38.03 (1) | 105.7 (4) |
| 60 | 3.73 (4) | 33.95 (4) | 91.23 (1) |
| 61 | 53.34 (4) | 27.58 (4) | 71.26 (4) |
| 62 | 90.72 (1) | 21.42 (4) | 93.1 (4) |
| 63 | -7.84 (4) | 30.1 (1) | 91.98 (4) |
| 64 | 0.84 (4) | 17.08 (4) | 116.2 (1) |
| 65 | 74.34 (4) | 44.38 (4) | 98.7 (1) |
| 66 | 6.3 (4) | 18.9 (4) | 86.38 (4) |
| 67 | 2.66 (1) | 24.78 (4) | 133.42 (4) |
| 68 | 16.88 (4) | 2.52 (1) | 129.78 (4) |

Table S3: The affinity values (kcal $mol^{-1}$) of the ligands (D-glucose, ONP and DDAO) calculated using AutoDock Vina, via Docking Pie, a Molecular Docking plugin for PyMOL into each of the 68 binding cavities of BSA.[8–10] n/a represents the binding cavities whereby the ligands were unable to successfully dock.

| Binding Cavity Sites | Ligand | | |
|---|---|---|---|
| | D-glucose | ONP | DDAO |
| 1 | -3.9 | -5.5 | -4.4 |
| 2 | n/a | -4.4 | -1.3 |
| 3 | -1.9 | n/a | -3.7 |
| 4 | -2.8 | -5.1 | n/a |
| 5 | -1.4 | -2.4 | -2.6 |
| 6 | n/a | n/a | -1.7 |
| 7 | -3.2 | -4.2 | -5.9 |
| 8 | -1.2 | -3.1 | -7.1 |
| 9 | -4.4 | -5.2 | n/a |
| 10 | n/a | n/a | -1.1 |
| 11 | -3.1 | -4.4 | -5.9 |
| 12 | -2.5 | -3.5 | -5.5 |
| 13 | -3.1 | -4.8 | n/a |
| 14 | n/a | -5.6 | -3.9 |
| 15 | -3.1 | -4 | -4.2 |
| 16 | -1.1 | -3.5 | -5 |
| 17 | -2.9 | -4 | -3.7 |
| 18 | n/a | n/a | n/a |
| 19 | -2.8 | -4.2 | -4.6 |
| 20 | -2.5 | -3.5 | -4.3 |
| 21 | -1.9 | -3.8 | -3.4 |
| 22 | n/a | n/a | -3.5 |
| 23 | -3.6 | -4.5 | -5.3 |
| 24 | n/a | -5.2 | -5.1 |
| 25 | -3.5 | -3.5 | -3.4 |
| 26 | -2.4 | -4.7 | -5.1 |
| 27 | -3.3 | -3.2 | -3.4 |
| 28 | -3.3 | -3.9 | n/a |
| 29 | 2.5 | -4.5 | -6.5 |
| 30 | -3.1 | -4 | -4.4 |
| 31 | 2.6 | -2.9 | -6.1 |
| 32 | n/a | n/a | -2.4 |
| 33 | -2.8 | -3 | -2.1 |
| 34 | -3.2 | -4.9 | -5 |
| 35 | -2.7 | -5.1 | -4.7 |
| 36 | n/a | n/a | n/a |
| 37 | 0.7 | n/a | -1.5 |
| 38 | -2.9 | -3.4 | -4.2 |
| 39 | -3.6 | -4.7 | -4.7 |
| 40 | -2.5 | -5.1 | n/a |
| 41 | -2.3 | -3.5 | -3.4 |
| 42 | n/a | n/a | -3.6 |
| 43 | -1.9 | -3.8 | -3.9 |
| 44 | -2.4 | -3.5 | -3.6 |
| 45 | -2.7 | -3.9 | n/a |
| 46 | n/a | -3.1 | -5.2 |
| 47 | 3.7 | -2.6 | -3.8 |
| 48 | -3.3 | n/a | -5.2 |
| 49 | -3 | -2.8 | n/a |
| 50 | -2.3 | -4.1 | -3.7 |
| 51 | -3.5 | -3.8 | -5.6 |
| 52 | -3.4 | n/a | -4.6 |
| 53 | n/a | -4.1 | -4.3 |
| 54 | -3.1 | -4.4 | -4.5 |
| 55 | 11.5 | 0.3 | 0.8 |
| 56 | -3.4 | -3.8 | n/a |
| 57 | n/a | n/a | -4.6 |
| 58 | -4 | -4.9 | -4.1 |
| 59 | -0.9 | -3.4 | -2.3 |
| 60 | 24.8 | 7.3 | 7.1 |
| 61 | n/a | 15 | 6.1 |
| 62 | n/a | n/a | n/a |
| 63 | -3.1 | -4.4 | -4.3 |
| 64 | 26.7 | 1.7 | 9.6 |
| 65 | -2.8 | -3.6 | -4.5 |
| 66 | 1.2 | n/a | n/a |
| 67 | -3.1 | -3.9 | -4.6 |
| 68 | -2.9 | -4 | -3.3 |

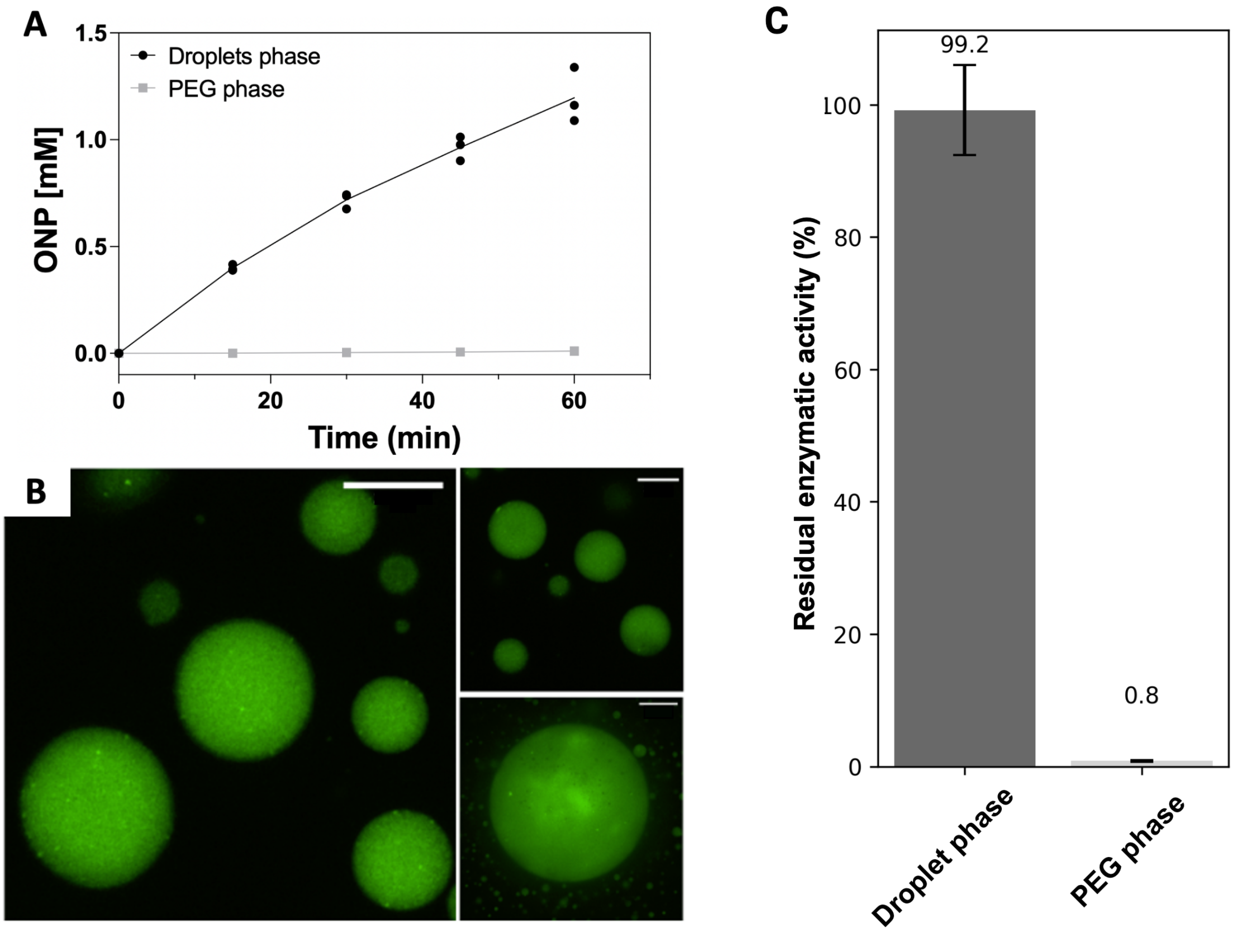

Figure S1: Evaluation of $\beta$-galactosidase partitioning inside the droplet phase. **A** Residual enzymatic activity in the PEG-rich (grey) and in the droplet phase (black) from an initial droplets mix containing 20 nM $\beta$-galactosidase in the presence of 5 mM ONP-gal. **B** Confocal microscope images of $\beta$-galactosidase labeled with Alexafluor-488 in the droplets. Scale bar 20 $\mu$m. **C** Residual enzymatic activity in the PEG-rich (grey) and in the droplet phase (black) calculated from the simple linear regression (y= $\beta$0+ $\beta$1x) of the data in panel A. The total enzymatic activity is defined as the sum of the slopes ($\beta$1) of the PEG and droplets phase. The data represent the mean ± SD.

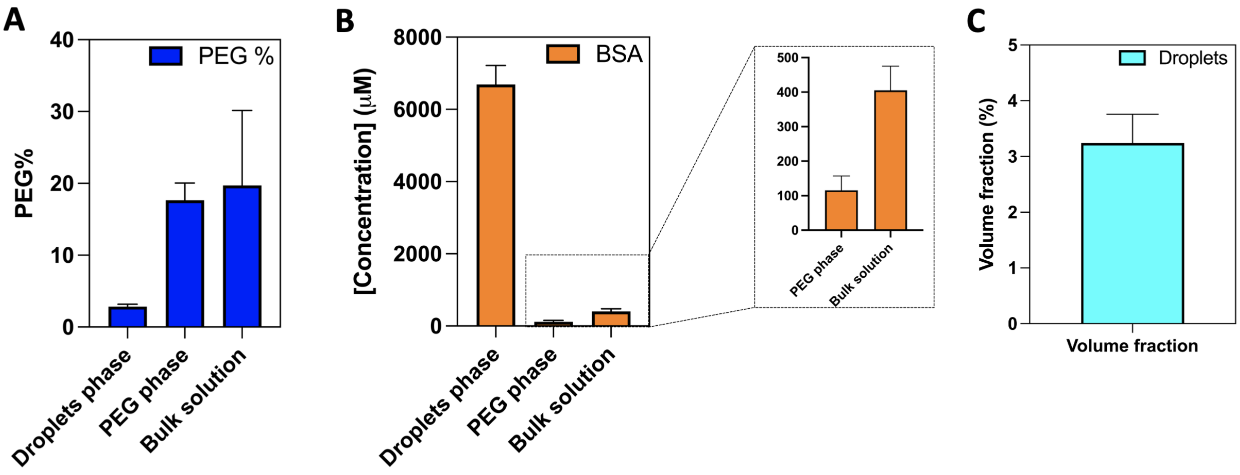

Figure S2: PEG-BSA system characterization. **A** PEG % (w/w) determination in the different liquid phases by NMR. 1 ml of droplets mixture was divided in two aliquots. Then, one aliquot was centrifuged (16000 rcf for 30 mins) to separate the droplet phase and the PEG phase. The PEG % (w/w) inside the droplet phase, the PEG phase and the bulk solution were obtained using NMR as previously reported.[1] The data represents the mean values ± standard deviation (SD) from four independent experiments (n≥4).**B** BSA concentration inside the droplets, PEG phase, and bulk solution. The concentration of BSA was determined using a spectrophotometer: the absorbance at 280 nm of the samples was used to get the BSA concentration using the Lambert-Beer equation. The extinction coefficient for BSA used was 43824 $M^{-1}cm^{-1}$ and was determined using ProtParam (https://web.expasy.org/protparam/). The data represents the mean values ± SD (n≥3). **C** The droplets volume fraction % (v/v) in the PEG-BSA system. 1 ml of droplets solution was prepared and centrifuged (16000 rcf for 30 minutes) to separate the droplet phase to the PEG phase. Then, the volume of the droplet phase was measured by pipetting. The data represents the mean values ± SD (n=3).

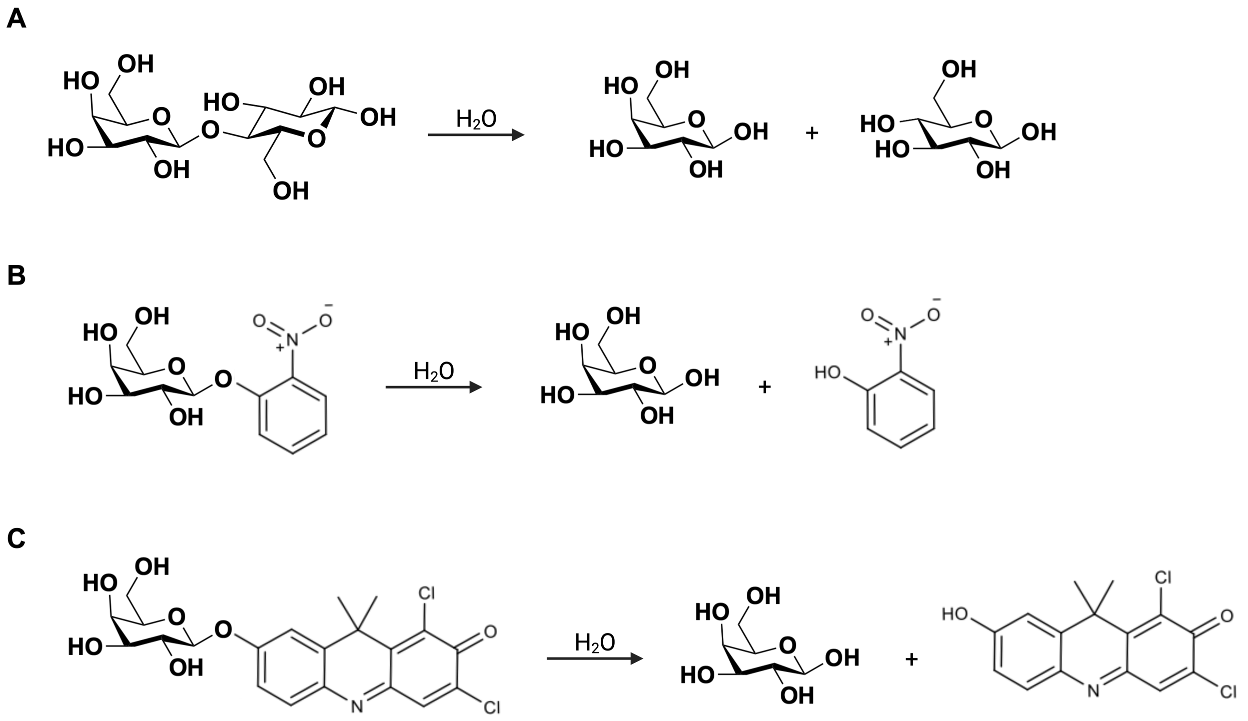

Figure S3: Schemes of the reactions catalyzed by the enzyme $\beta$-galactosidase. Scheme of the chemical reactions performed by $\beta$-galactosidase on the substrates D-lactose (**A**), ONP-gal (**B**), and DDAO-gal (**C**).

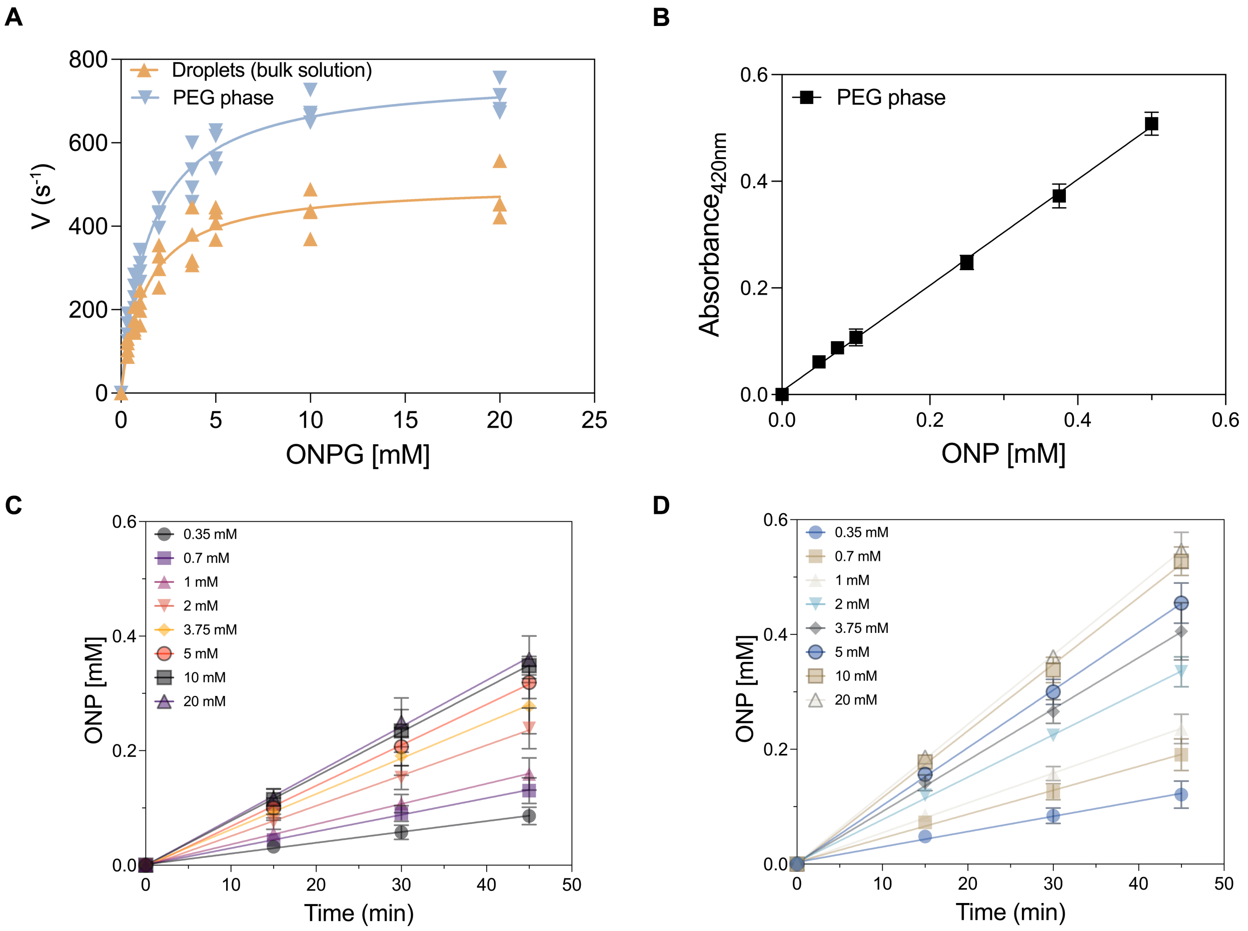

Figure S4: Michaelis-Menten curves of $\beta$-galactosidase using the substrate 4-nitrophenyl $\beta$-D-galactopyranoside (ONP-gal) **A** $\beta$-galactosidase Michaelis-Menten curve with an ONP-gal concentration range of 0.35 - 20 mM. The velocity (V) measurements were performed in the bulk solution (orange) and in PEG phase only (light blue). For the activity measurements in the bulk solution, the concentration of $\beta$-galactosidase in the droplets was 6 $\mu$M and the concentration of the enzyme in the overall solution was 0.286 nM during the kinetic assay. For the activity measurements in the PEG phase, the enzyme concentration was 0.286 nM during the kinetic assay. Data are represented as Michaelis-Menten curve fitting using Prism software, all data points are shown (n=4). **B** Calibration curve of ortho-nitrophenol (ONP) in the PEG phase. The calibration curve was used to convert the absorbance value into ONP concentration during the enzyme activity measurements. The data are represented as mean values $\pm$ SD ($n \geq 3$). **C** Linear trends of ONP production at several ONP-gal concentrations (0.35-20 mM) over time using 6 $\mu$M $\beta$-galactosidase partitioned in the droplets (0.286 nM $\beta$-galactosidase in the bulk solution). The data are represented as mean values $\pm$ SD ($n = 4$). **D** Linear trends of ONP production at several ONP-gal concentrations (0.35-20 mM) over time using 0.286 nM $\beta$-galactosidase in the PEG phase. The data are represented as mean values $\pm$ SD ($n = 4$).

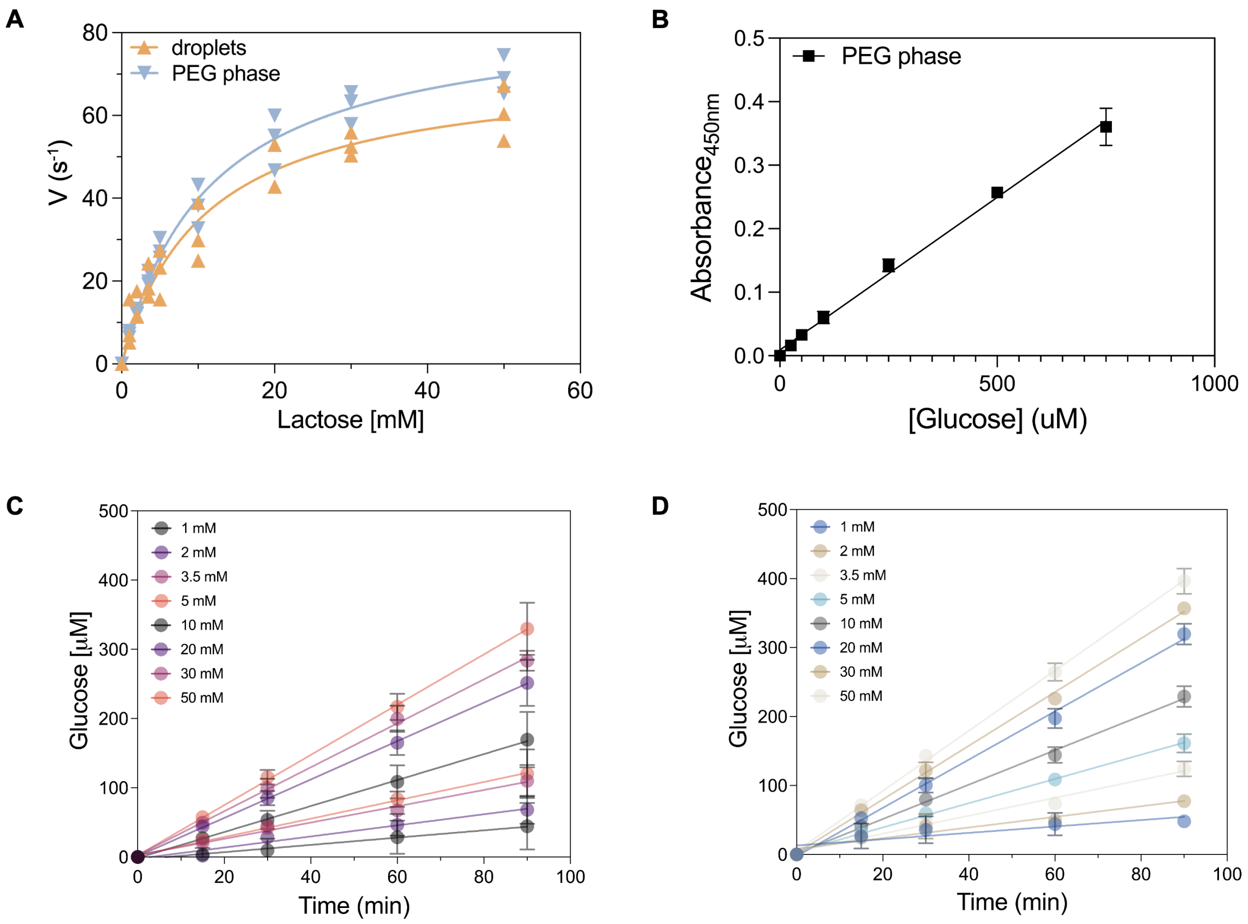

Figure S5: Michaelis-Menten curves of $\beta$-galactosidase using the substrate D-lactose **A** $\beta$-galactosidase Michaelis-Menten curve at different D-lactose concentrations (0.5 - 50 mM). The velocity (V) measurements were performed in the bulk solution (orange) and in PEG phase only (light blue). For the activity measurements in the droplet phase, the concentration of $\beta$-galactosidase in the droplets was 3 $\mu$M and the concentration of the enzyme in the bulk solution was 1 nM during the kinetic assay. For the activity measurements in the PEG phase, the enzyme concentration was 1 nM during the kinetic assay. Data are represented as Michaelis-Menten curve fitting using Prism software, all data points are shown (n=3). **B** Calibration curve of D-glucose in the PEG phase. The calibration curve was used to convert the absorbance value into D-glucose concentration during the enzyme activity measurements. The data are represented as mean values $\pm$ SD (n = 3).**C** Linear trends of D-glucose production at several lactose concentrations (1-50 mM) over time using 3 $\mu$M $\beta$-galactosidase partitioned in the droplets (1 nM $\beta$-galactosidase in the bulk solution). The data are represented as mean values $\pm$ SD (n = 3). **D** Linear trends of D-glucose production at several lactose concentrations (1-50 mM) over time using 1 nM $\beta$-galactosidase in the PEG phase. The data are represented as mean values $\pm$ SD (n = 3).

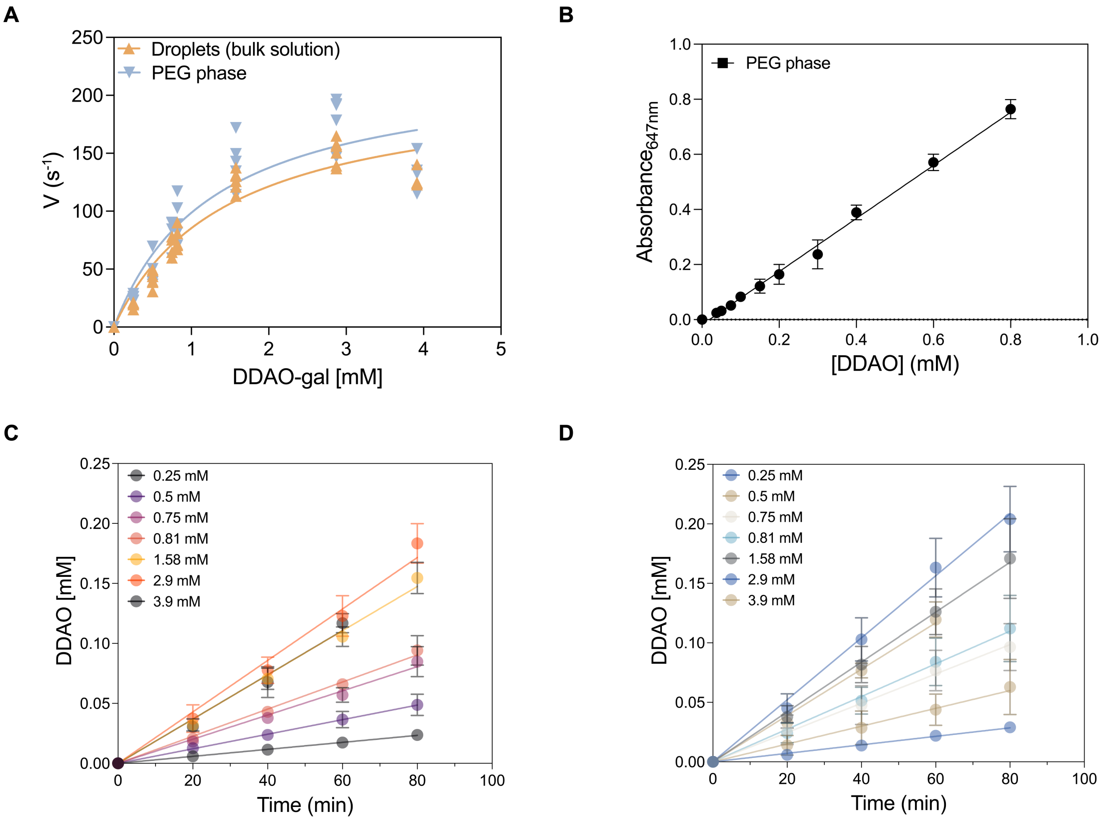

Figure S6: Michaelis-Menten curves of $\beta$-galactosidase using the substrate 9H-1,3-Dichloro-9,9-Dimethylacridin-2-One-7-yl $\beta$-D-galactopyranoside (DDAO-gal). **A** $\beta$-galactosidase Michaelis-Menten curve at different DDAO-gal concentrations (0.25-3.9 mM). The DDAO-gal concentration were normalized by the precipitated fraction (Fig.S20). The velocity (V) measurements were performed in the bulk solution (orange) and in PEG phase only (light blue). For the activity measurements in the bulk solution, the concentration of $\beta$-galactosidase in the droplets was 3 $\mu$M and the concentration of the enzyme in the bulk solution was 0.25 nM during the kinetic assay. For the activity measurements in the PEG phase, the enzyme concentration was 0.25 nM during the kinetic assay. Data are represented as Michaelis-Menten curve fitting using Prism software, all data points are shown ($n \geq 4$). **B** Calibration curve of 7-hydroxy-9H-1,3-dichloro-9,9-dimethylacridin-2-one (DDAO) in the PEG-rich phase. The calibration curve was used to convert the absorbance value into DDAO concentration during the enzyme activity measurements. The data are represented as mean values $\pm$ SD ($n \geq 3$).**C** Linear trends of DDAO production at different DDAO-gal concentrations (0.25-3.5 mM) over time using 3 $\mu$M $\beta$-galactosidase partitioned in the droplets (0.25 nM $\beta$-galactosidase in the bulk solution). The data are represented as mean values $\pm$ SD ($n \geq 3$). **D** Linear trends of DDAO production at several DDAO-gal concentrations (0.25-3.5 mM) over time using 0.25 nM $\beta$-galactosidase in the PEG phase. The data are represented as mean values $\pm$ SD ($n = 3$).

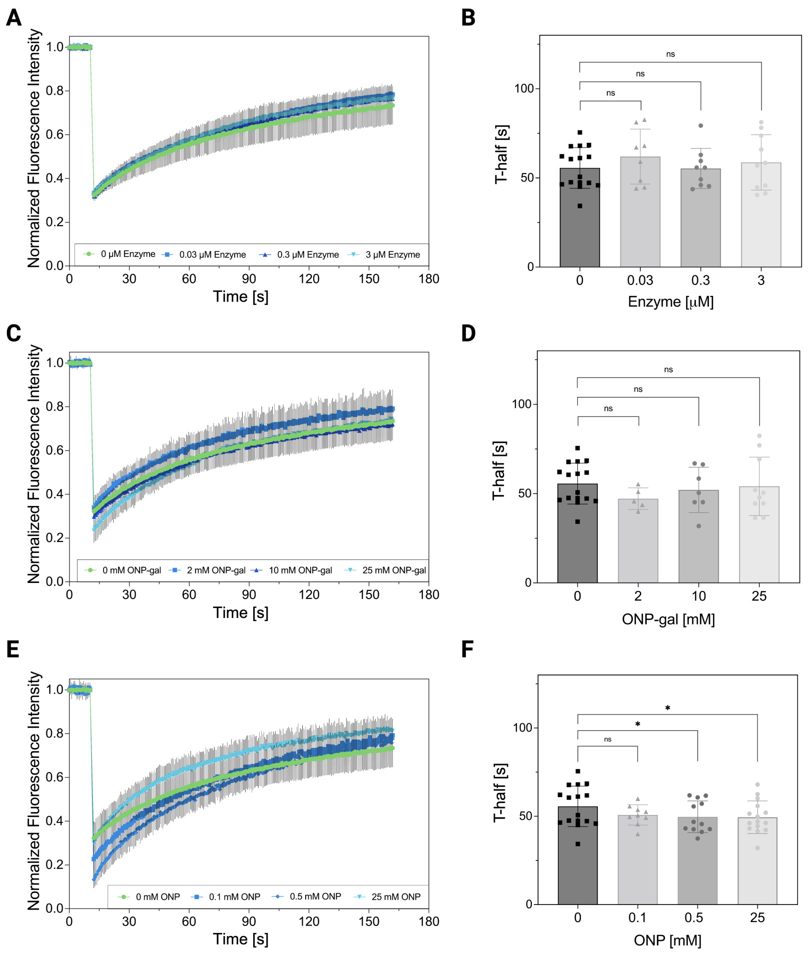

Figure S7: Negative controls of FRAP experiments in the presence of the reaction substrate, product and enzyme. **A** Normalized FRAP data following the photobleaching of (Alexafluor488-labeled) BSA droplets in the presence of $\beta$-galactosidase (0-3$\mu$M inside the droplets). The data represent the mean ± SD (n≥8). **B** Relative FRAP half-recovery time of droplets containing a range of $0-100\ nM$ enzyme in the solution ($0-3\ \mu$M $\beta$-galactosidase inside the droplets). The boxes represent the mean values and the whiskers are the SD (n≥8). **C** Normalized FRAP data following the photobleaching of (Alexafluor488-labeled) BSA droplets in the presence of the reaction substrate (0-25 mM ONP-gal). The data represent the mean ± SD (n≥5). **D** Relative FRAP half-recovery time of droplets containing a range of the reaction substrate (0-25 mM ONP-gal). The boxes represent the mean values and the whiskers are the SD (n≥5). **E** Normalized FRAP data following the photobleaching of (Alexafluor488-labeled) BSA droplets in the presence of the reaction product (0-25 mM ONP). The data represent the mean ± SD (n≥14). **F** Relative FRAP half-recovery time of droplets containing a range of the reaction product (0-25 mM ONP). The boxes represent the mean values and the whiskers are the SD (n≥8). $*P < 0.05$, $**P < 0.01$, $***P < 0.001$.

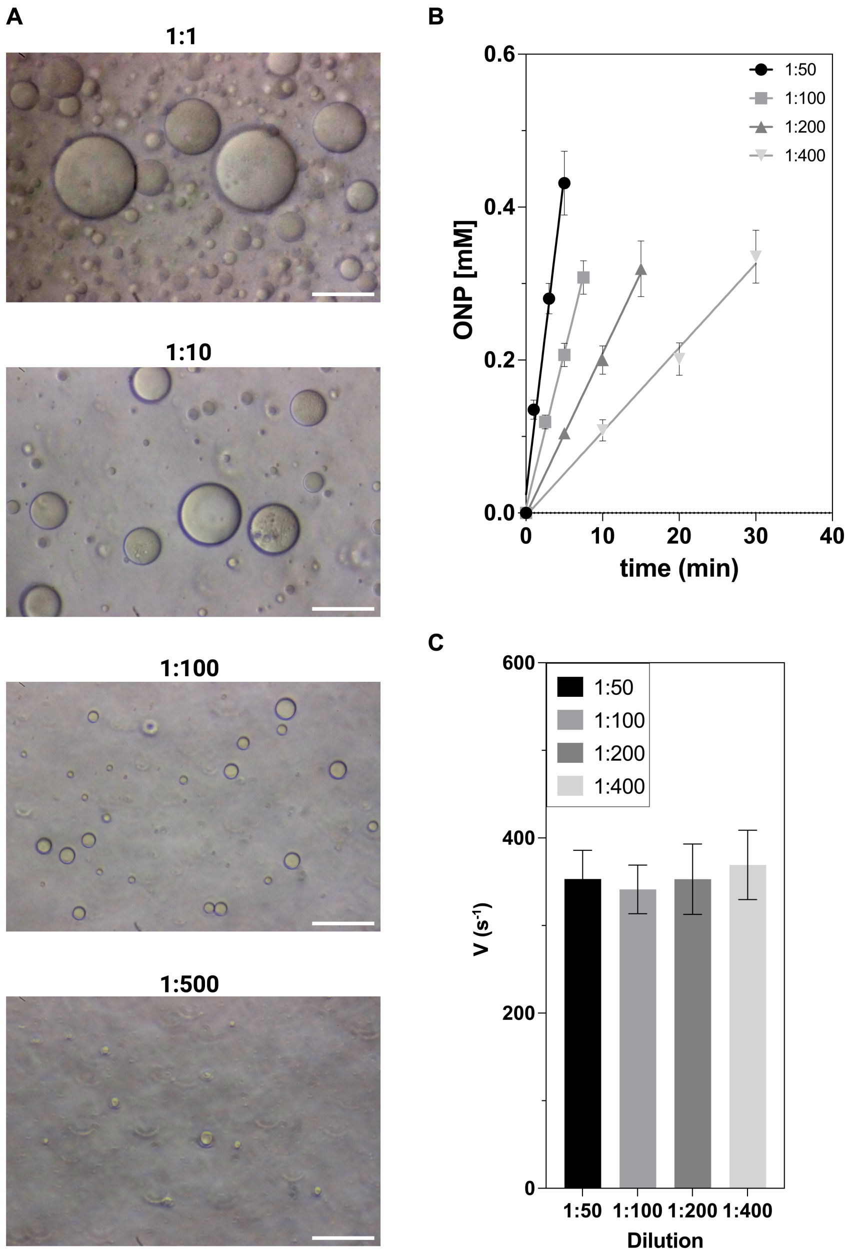

Figure S8: Dilution decreases droplets size and avoids substrate diffusion limitation. **A** Light microscope images (Olympus CKX41, 40x magnification lens) of the droplets solution diluted 1:1, 1:10, 1:100 and 1:500. The droplets solution was diluted using PEG-rich phase solution. Scale bar 50 $\mu$m. **B** Linear trends of ONP production at different droplets dilution (1:50, 1:100, 1:200 and 1:400) over time using 6 $\mu$M $\beta$-galactosidase partitioned in the droplets in the presence of 25 mM ONP-gal. The data are represented as mean values $\pm$ SD (n $\geq$ 3). **C** Catalytic rate of the enzymes is not affected in the tested droplet dilution. The slope from Panel B was converted into enzyme catalytic velocity ($s^{-1}$) by normalizing it for the final enzyme concentration. The data represents the mean values $\pm$ SD (n$\geq$ 3).

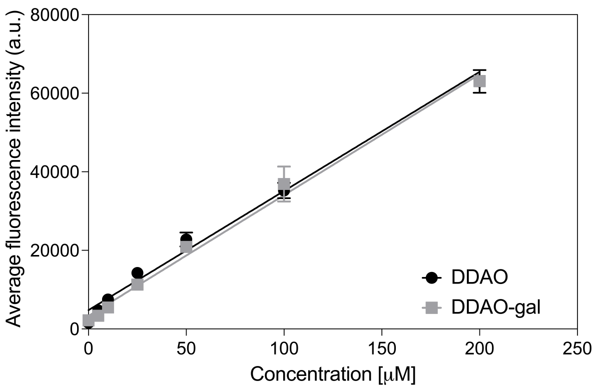

Figure S9: Calibration curve of DDAO and DDAO-gal for the confocal microscope experiments. The dataset represents the average intensity of droplets over a range of concentrations of the reaction substrate (DDAO-gal, 0-200 $\mu$M) and product (DDAO, 0-200 $\mu$M). The fluorescence intensity was measured on droplets of a diameter range of 50-500 $\mu$m after two hours from the addition of DDAO/DDAO-gal. The dataset was recorded with the confocal microscope LSM880 (20x lens) by exciting the probes at $\lambda_g$ = 488 nm and $\lambda_g$ = 646 nm and by recording the signal at the range of $\lambda_g$ = 588-624 nm and $\lambda_g$ = 657-680 nm for DDAO-gal and DDAO, respectively. The data represents the mean values $\pm$ SD (n$\geq$6). The slopes of the linear fitting were used to build the calibration curve in the Mathlab script to analyze the diffusion of DDAO-gal and DDAO.

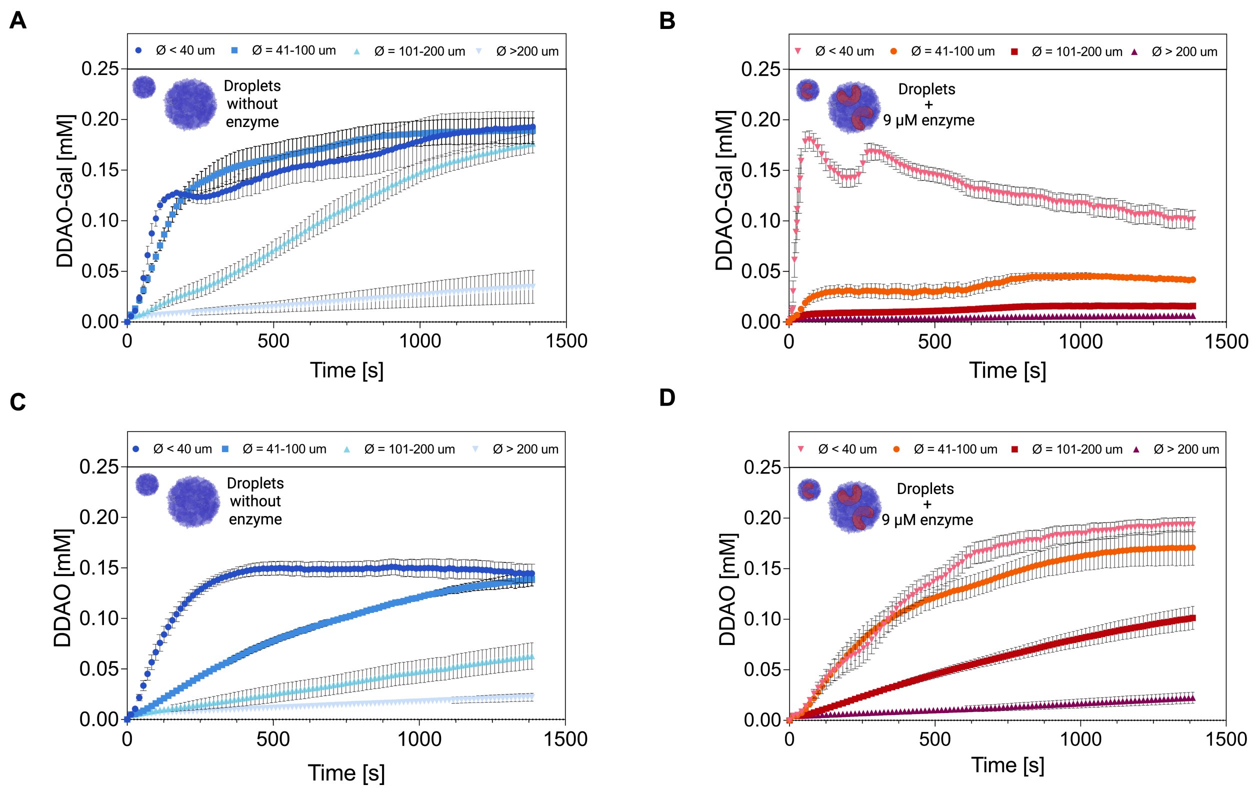

Figure S10: Diffusion/generation of DDAO and DDAO-gal inside the BSA droplets in the presence and absence of $\beta$-galactosidase. **A** Diffusion of the reaction substrate DDAO-gal inside the BSA droplets. The plots show the average concentration of DDAO-gal over time inside the droplets of different radius ranges. The data are obtained with a Matlab script from confocal microscope movies (n=9) recording the droplets for 1500 seconds after the addition of 0.2 mM DDAO-gal. The data represents the mean values ± SEM (n≥4). **B** Diffusion of the reaction substrate DDAO-gal inside enzyme containing (9$\mu$M) droplets. The plots show the average concentration of DDAO-gal over time inside the droplets of different radius ranges. The data are obtained with a Matlab script from confocal microscope videos (n=10) recording the droplets for 1500 seconds after the addition of 0.2 mM DDAO-gal. The data represents the mean values ± SEM (n≥4). **C** Diffusion of the reaction product DDAO inside the droplets. The plots show the average concentration of DDAO over time inside the droplets of different radius ranges. The data are obtained with a Matlab script from confocal microscope movies (n=7) recording the droplets for 1500 seconds after the addition of 0.2 mM DDAO. The data represents the mean values ± SEM (n≥3). **D** Generation of the reaction product DDAO inside enzyme containing (9$\mu$M) BSA droplets. The plots show the average concentration of DDAO inside the droplets of different radius ranges. The data are obtained with a Matlab script from confocal microscope videos (n=10) recording the droplets for 1500 seconds after the addition of 0.2 mM DDAO-gal. The data represents the mean values ± SEM (n≥4).

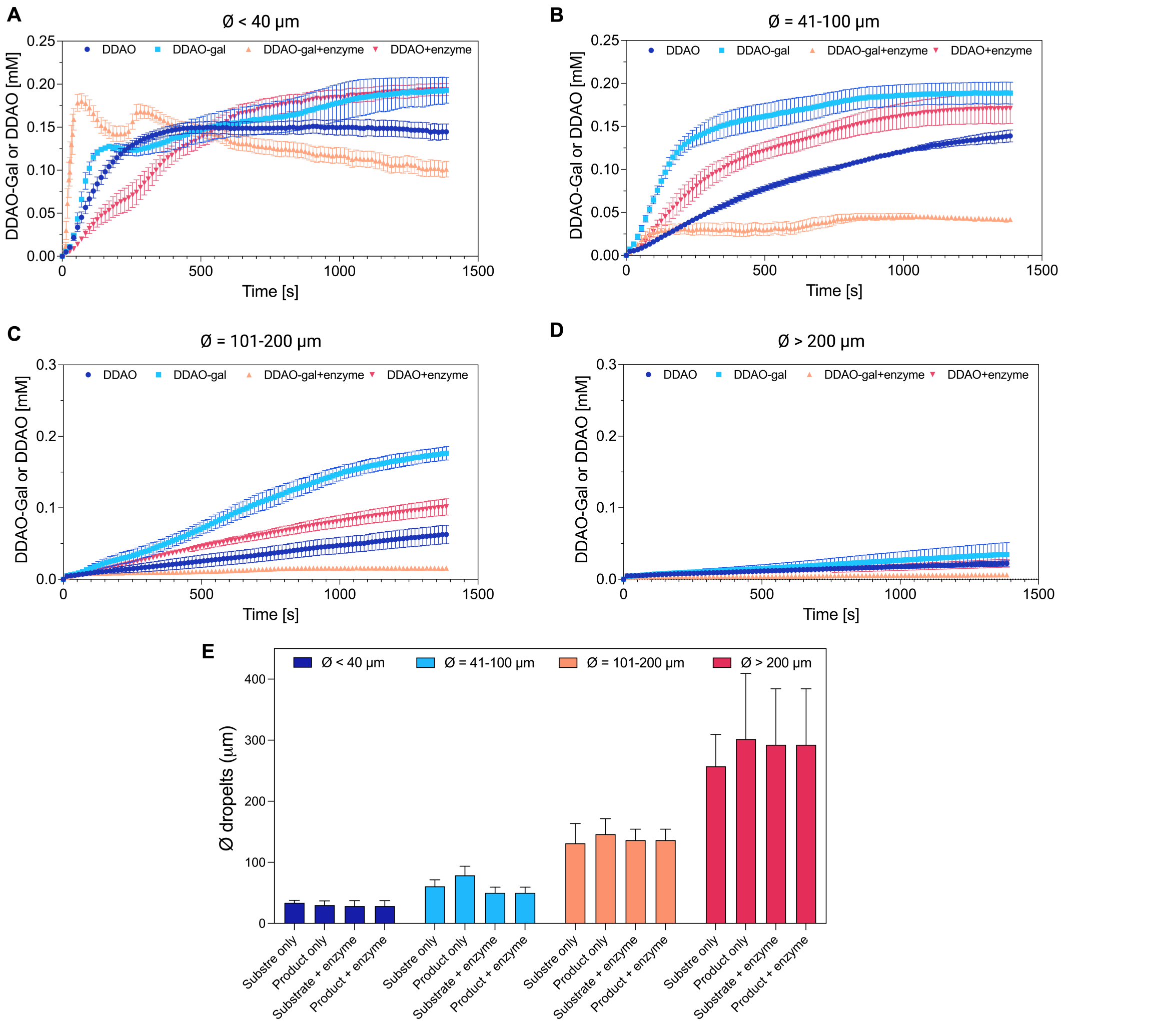

Figure S11: Comparison in droplets of different sizes of the diffusion/generation of DDAO-gal and DDAO inside the BSA droplets in the presence and absence of $\beta$-galactosidase. **A–D** Average concentration of DDAO-gal and DDAO over time inside BSA droplets of varying diameters: **A** ¡40$\mu$m, **B** 41–100$\mu$m, **C** 101–200$\mu$m, **D** ¿201$\mu$m. The plots show the diffusion of DDAO-gal and DDAO in the presence and absence of the enzyme (9 $\mu$M). The data are obtained with a Matlab script from confocal microscope movies (n$\geq$4) recorded for 1500s after addition of 0.2mM DDAO-gal or DDAO. The data represents the mean values $\pm$ SEM (n$\geq$4). **E** Average droplet size of each sample group. The histograms represent the mean $\pm$ SD (n$\geq$4).

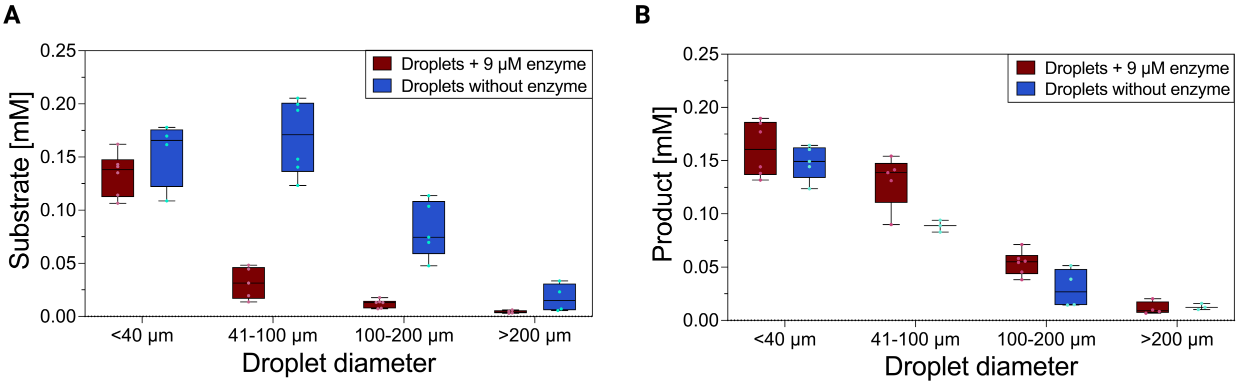

Figure S12: Substrate and Product concentration inside the droplets 10 minutes of the addition of the substrate/product. **A** Substrate concentration inside the droplets after 10 minutes from the addition of 0.2 mM DDAO-Gal in droplets containing 9 $\mu$M $\beta$-galactosidase (red) and without enzyme (blue). The analysis was performed on droplets of a $\varnothing$ range of 15-398$\mu$m (n$\geq$3). The central mark indicates the median, the top and bottom edges of the box the 25th and 75th percentile, and the whiskers the maximum and minimum value of the data. **B** Product concentration inside the droplets after 10 minutes from the addition of 0.2mM DDAO-galactoside in droplets containing 9 $\mu$M $\beta$-galactosidase (red) and without enzyme (blue). The analysis was performed on droplets of $\varnothing$ range of 15-426$\mu$m (n$\geq$3). The central mark indicates the median, the top and bottom edges of the box the 25th and 75th percentile, the whiskers the maximum and minimum value of the data.

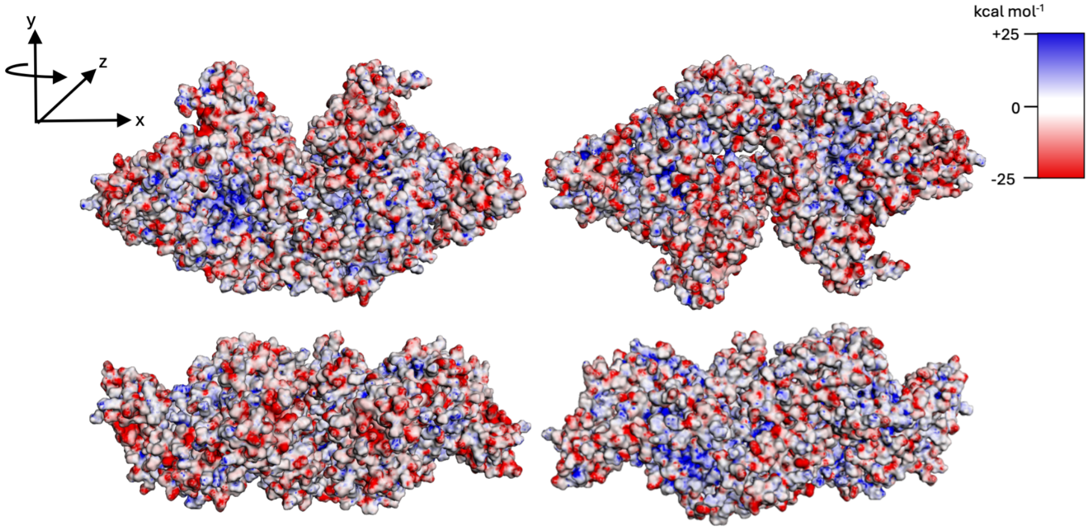

Figure S13: BSA electrostatic charge analysis. Electrostatic potential (blue positive and red negative) map of the protein surface calculated using Adaptive Poisson-Boltzmann Solver (APBS).[11]

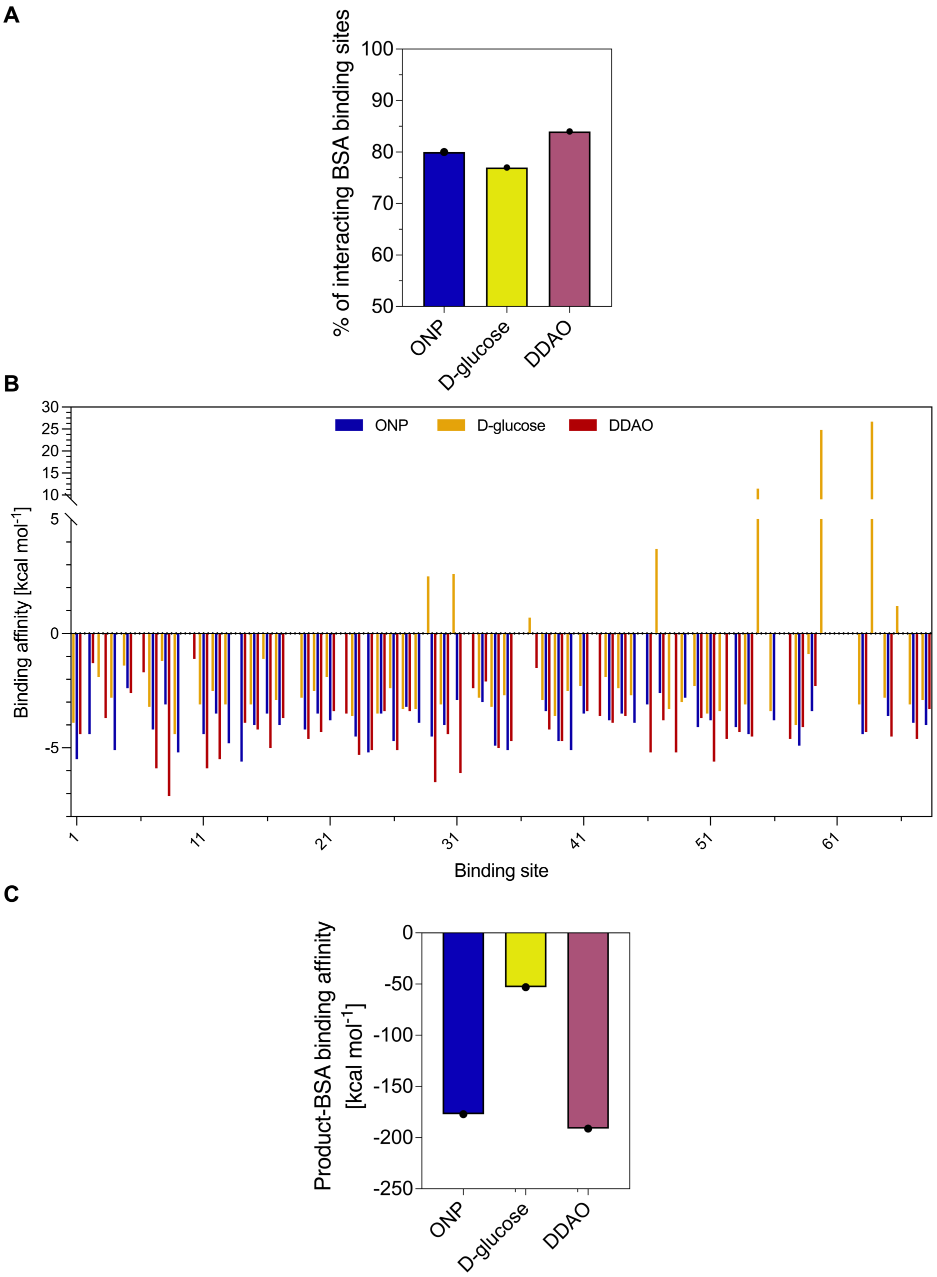

Figure S14: Reaction product binding analysis with BSA cavities. **A** Percentage of the total 68 BSA binding cavities interacting with each reaction product. **B** The affinity values (kcal mol$^{-1}$) of the ligands (D-glucose, ONP and DDAO) into each of the 68 binding cavities of BSA calculated using AutoDock Vina, via Docking Pie, a Molecular Docking plugin for PyMOL (PyMOL Molecular Graphics System, Version 3.0 Schrödinger, LLC.).[8–10] **C** The overall affinity values (kcal mol-1) of the ligands (D-glucose, ONP and DDAO) docking in the protein surface cavities calculated using AutoDock Vina, via Docking Pie, a Molecular Docking plugin for PyMOL.

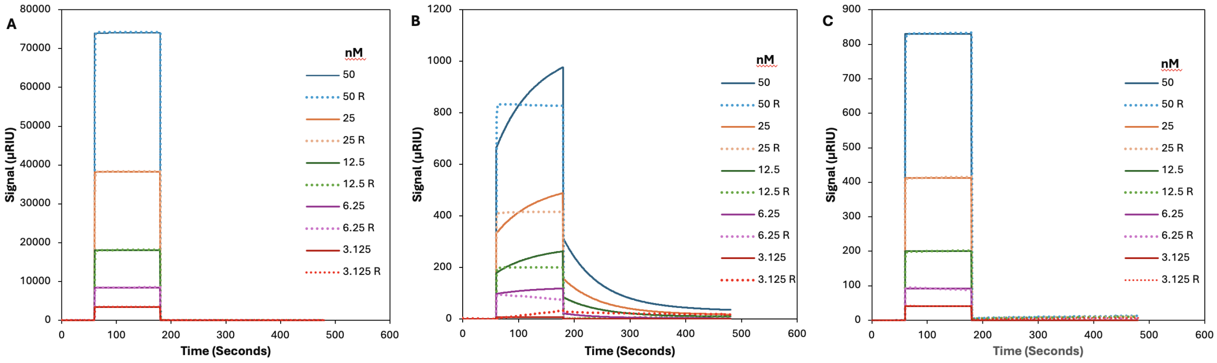

Figure S15: Representative fitted sensorgrams (solid line) with corresponding raw data (dotted line) of the molecular interactions of BSA immobilised onto a PEG coated gold SPR chips for five concentrations (nM) of the reactions products with: **A** ONP binding to BSA **B** DDAO binding to BSA **C** D-glucose binding to BSA.

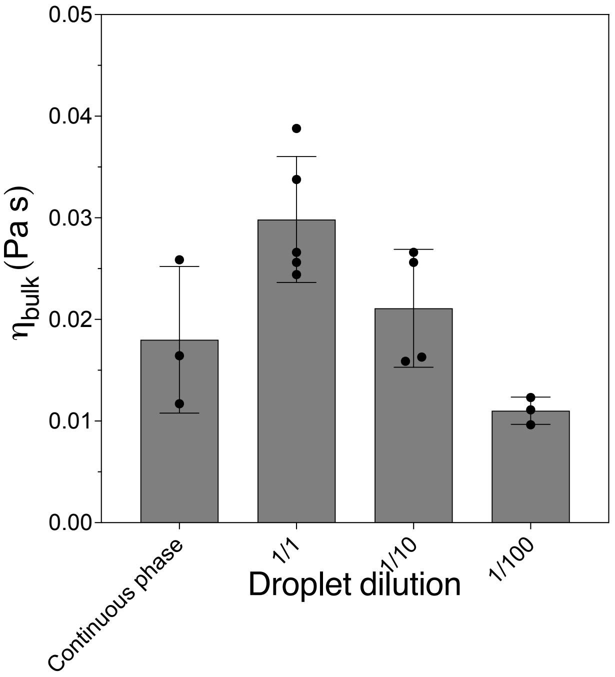

Figure S16: Effect of droplet volume fraction on bulk viscosity ($\eta_{\mathrm{bulk}}$). To evaluate how the concentration of the droplet phase influences $\eta_{\mathrm{bulk}}$, droplet mixtures were measured at different dilutions with the PEG-rich phase. Dilution ratios of 1/1, 1/10, 1/100 and continuous-phase only correspond to droplet volume fractions of approximately 3%, 0.3%, and 0.03% and 0%, respectively. The data represent the mean values ± SD (n $\geq$ 3).

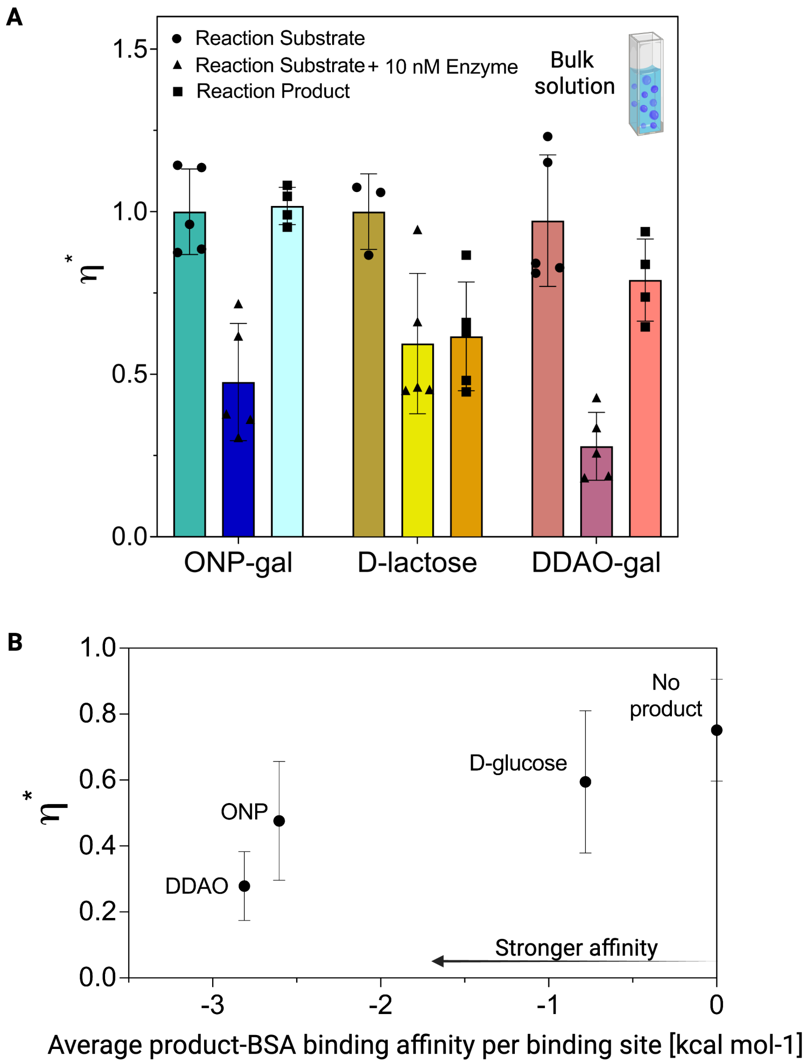

Figure S17: Bulk rheology measurements in presence and absence of enzymatic activity. **A** Relative viscosity ($\eta^*$) of the PEG solvent–BSA droplet bulk solution under three conditions: (i) substrate in the PEG-rich solvent without enzyme (circles), (ii) substrate in the PEG-rich solvent with enzyme localized in BSA-rich droplets (triangles), and (iii) product in the PEG-rich solvent, no enzyme in the BSA droplets (squares). Enzymatic activity (10 nM $\beta$-galactosidase in bulk) leads to a significant decrease in ($\eta^*$), especially for ONP-gal and DDAO-gal, suggesting that substrate–BSA interactions are altered upon catalysis. Product conditions show a partial rebound in ($\eta^*$), particularly for substrates with stronger BSA affinity. Data represent mean ± SD ($n \geq 3$). **B** Comparison of the activity-induced relative viscosity change ($\eta^*$) with the average product–BSA binding affinity determined from molecular docking (Fig. 4B). The three reaction systems exhibit a similar ordering, with stronger product–BSA interactions associated with greater activity-dependent viscosity modulation. Data represent mean ± SD ($n \geq 4$).

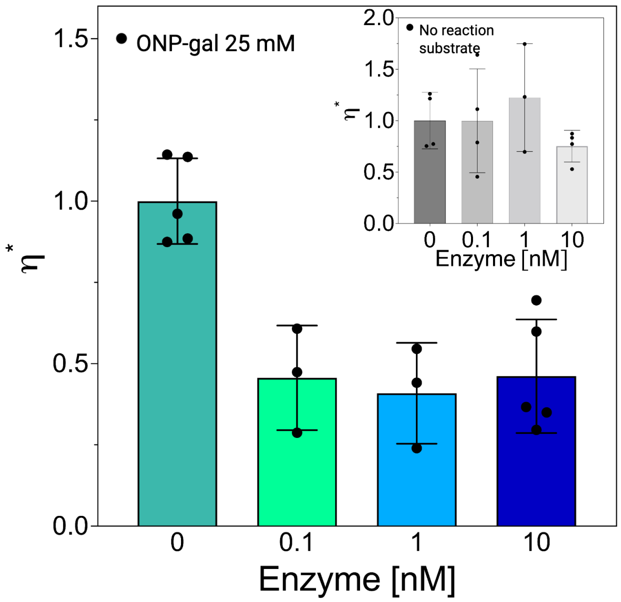

Figure S18: Bulk rheology measurements of the bulk solution (continuous phase + droplets) with different enzyme concentrations. Relative viscosity ($\eta^*$) obtained at different $\beta$-galactosidase concentration (0, 0.1, 1, and 10 nM) in the presence of 25 mM ONP-gal. A decreased $\eta^*$ is always associated with the presence of active enzyme. The inset shows that varying enzyme concentrations do not affect the shear viscosity of the solution in the absence of substrate. The data represents the mean values ± SD (n≥3).

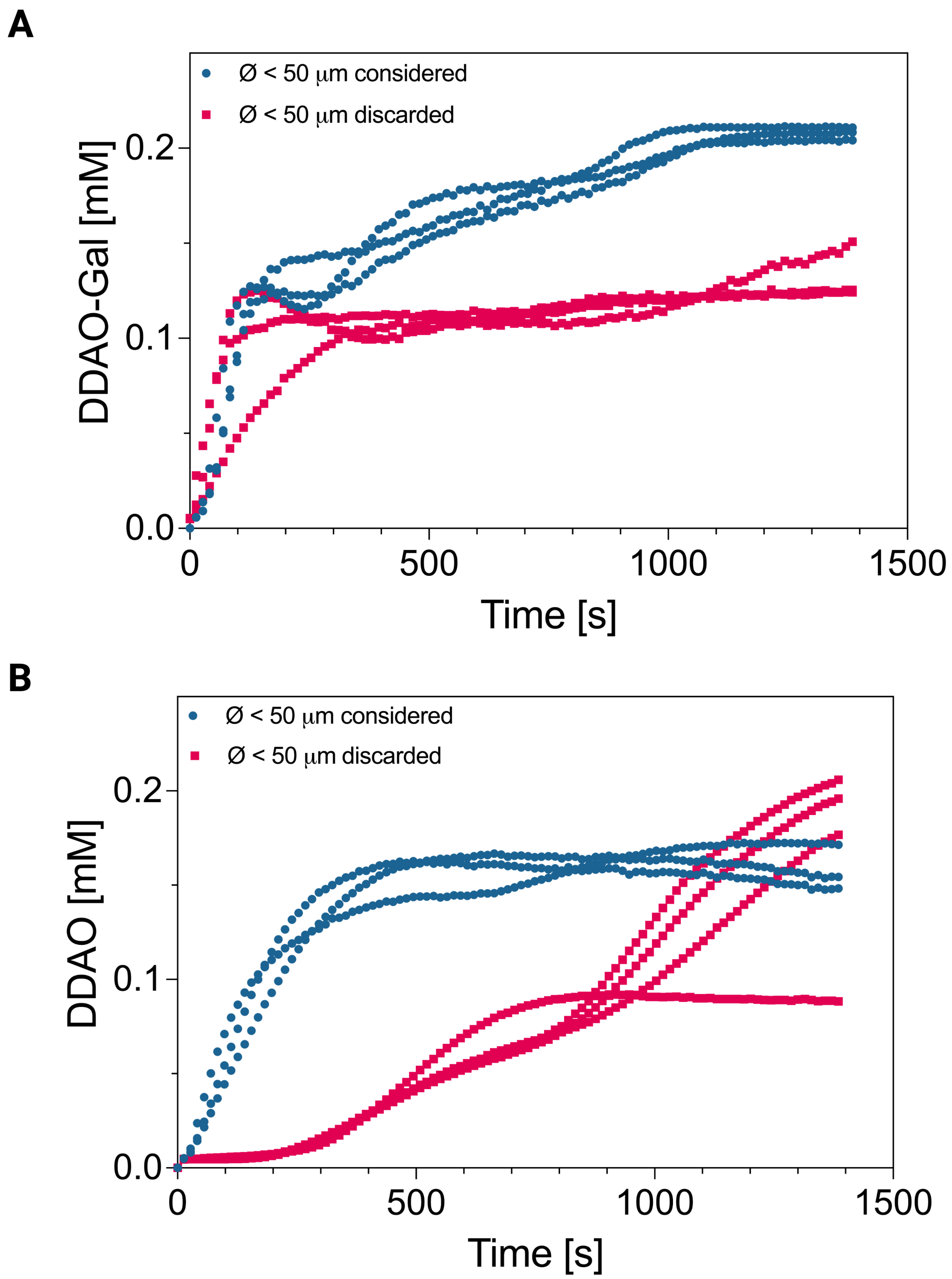

Figure S19: Filtering of the diffusion patterns for the diffusion analysis. **A** Representative diffusion patterns of DDAO-gal (0.2 mM) in BSA-only droplets. The plot highlights the diffusion patterns considered (blue) and discarded (red) for the diffusion analysis. The reported experiments were discarded because the concentration of substrate did not reach the concentration of substrate added. **B** Representative non-linear diffusion patterns of DDAO (0.2 mM) in BSA droplets without enzyme. The plot shows examples of the diffusion patterns considered (blue) and discarded (red) for the diffusion analysis. The latter experiments were discarded because they showed non-linear or delayed diffusion compared to the considered ones.

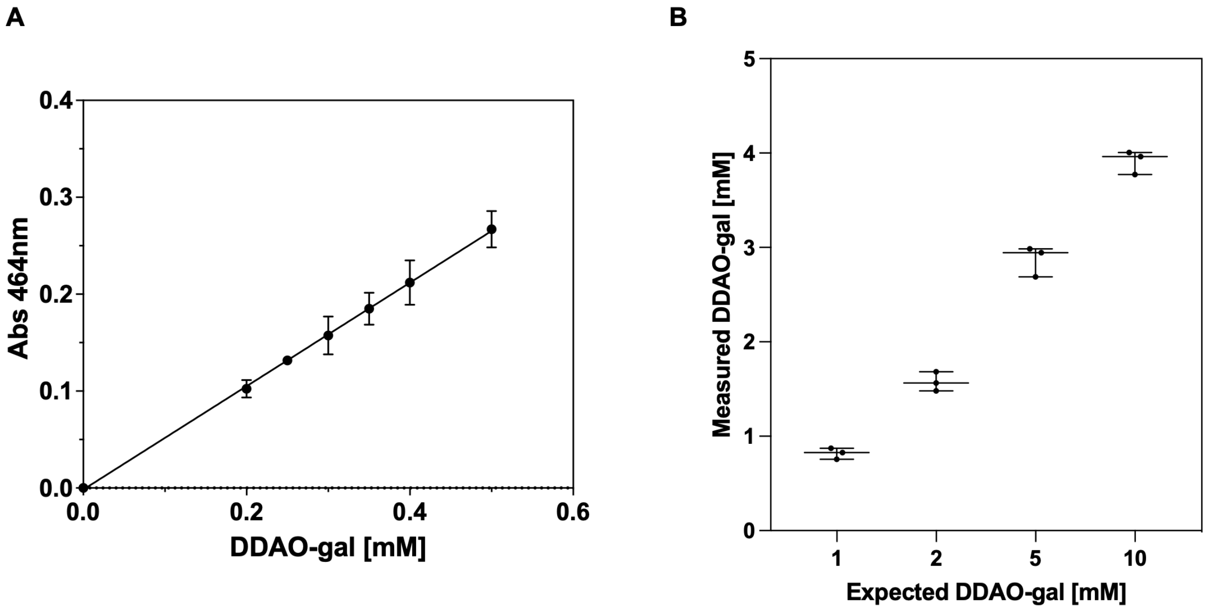

Figure S20: DDAO-gal precipitated fraction determination. **A** Calibration curve of DDAO-gal recorded using 5 $\mu$L of standard solution into 250 $\mu$L of TRIS 0.3 M, pH 7.0 and by measuring the absorbance at 464 nm. The data represents the mean values ± SD (n=3). **B** The DDAO-gal samples that showed precipitation were centrifuged and the actual concentration of the samples were determined by recording their absorbance at 464 nm and by using the slope of the calibration curve (A) to determine their concentration. The horizontal line represents the mean values ± SD, all data points are shown (n=3).

#### Supplementary movies

- **Supplementary movie 1.** Confocal microscopy movie of red fluorescent particle localized inside the enzyme-less droplets in the presence of 25 mM ONP-gal. The movie was recorded using a bright field and a $\lambda_g$= 561 nm laser channel at 2.12 fps. The channels were merged and the intensity was adjusted using Fiji ImageJ software. The movie represent a 2 minutes recording, the fps was sped up 10 times (21 fps). scale bar 25 $\mu$m.

- **Supplementary movie 2.** Confocal microscopy movie of red fluorescent particle localized inside the droplets containig 0.03 $\mu$M $\beta$-galactosidase in the presence of 25 mM ONP-gal. The movie was recorded using a bright field and a $\lambda_g$= 561 nm laser channel at 2.12 fps. The channels were merged and the intensity was adjusted using Fiji ImageJ software. The movie fps was sped up 10 times (21 fps). The screenshots shown in Fig.3A were obtained from this movie. Scale bar 25 $\mu$m.

- **Supplementary movie 3.** Confocal microscopy movie of red fluorescent particle localized inside the droplets containig 0.3 $\mu$M $\beta$-galactosidase in the presence of 25 mM ONP-gal. The movie was recorded using a bright field and a $\lambda_g$= 561 nm laser channel at 2.12 fps. The channels were merged and the intensity was adjusted using Fiji ImageJ software. The movie represent a 2 minutes recording, the fps was sped up 10 times (21 fps). Scale bar 25 $\mu$m.

- **Supplementary movie 4.** Confocal microscopy movie of red fluorescent particle localized inside the droplets containig 3 $\mu$M $\beta$-galactosidase in the presence of 25 mM ONP-gal. The movie was recorded using a bright field and a $\lambda_g$= 561 nm laser channel at 2.12 fps. The channels were merged and the intensity was adjusted using Fiji ImageJ software. The movie represent a 2 minutes recording, the fps was sped up 10 times (21 fps). Scale bar 25 $\mu$m.

- **Supplementary movie 5.** Confocal microscopy movie of FRAP experiment on droplet without enzyme in the presence of 25 mM ONP-gal. The droplets containing 15% Alexafluor 488-BSA were subjected to the FRAP experiment using a $\lambda_g$= 488 nm laser, then the fluorescence recovery was measured for 120 seconds at 1.83 fps. The movie fps have been modified to convert the video timescale to 0.5 minute per second. Scale bar 20 $\mu$m.

- **Supplementary movie 6.** Confocal microscopy movie of FRAP experiment on droplet containing 0.03$\mu$M $\beta$-galactosidase in the presence of 25 mM ONP-gal. The droplets containing 15% Alexafluor 488-BSA were subjected to the FRAP experiment using a $\lambda_g$= 488 nm laser, then the fluorescence recovery was measured for 120 seconds at 1.83 fps. The screenshots shown in Fig.3A were obtained from this movie. The movie fps have been modified to convert the video timescale to 0.5 minute per second. Scale bar 20 $\mu$m.

- **Supplementary movie 7.** Confocal microscopy movie of FRAP experiment on droplet containing 0.3$\mu$M $\beta$-galactosidase in the presence of 25 mM ONP-gal. The droplets were subjected to the FRAP experiment, then the fluorescence recovery was measured for 120 seconds at 1.83 fps. The movie fps have been modified to convert the video timescale to 0.5 minute per second. Scale bar 20 $\mu$m.

- **Supplementary movie 8.** Confocal microscopy movie of FRAP experiment on droplet containing 3$\mu$M $\beta$-galactosidase in the presence of 25 mM ONP-gal. The droplets were subjected to the FRAP experiment, then the fluorescence recovery was measured for 120 seconds at 1.83 fps. . The movie fps have been modified to convert the video timescale to 0.5 minute per second. Scale bar 20 $\mu$m.

- **Supplementary movie 9.** Diffusion of DDAO-gal and generation of the reaction product DAAO inside droplets containing 9$\mu$M $\beta$-galactosidase. The enzyme-BSA droplets are shown in the bright field channel, while the diffusion of DDAO-gal and the generation of the reaction product DDAO are displayed in the green ($\lambda_g = 488$ nm) and red channels ($\lambda_g = 648$ nm), respectively. The movie was recorded for 8 minutes at 0.94 fps. The movie fps have been modified to convert the video timescale to 1 minute per second. The scale bar is 100 $\mu$m.
